# Transcription-induced active forces suppress chromatin motion

**DOI:** 10.1101/2022.04.30.490180

**Authors:** Sucheol Shin, Guang Shi, Hyun Woo Cho, D. Thirumalai

## Abstract

The organization of interphase chromosomes in a number of species is starting to emerge thanks to advances in a variety of experimental techniques. However, much less is known about the dynamics, especially in the functional states of chromatin. Some experiments have shown that the mobility of individual loci in human interphase chromosome decreases during transcription, and increases upon inhibiting transcription. This is a counter-intuitive finding because it is thought that the active mechanical force (*F*) on the order of ten pico-newtons, generated by RNA polymerase II (RNAPII) that is presumably transmitted to the gene-rich region of the chromatin, would render it more open, thus enhancing the mobility. Inspired by these observations, we developed a minimal active copolymer model for interphase chromosomes to investigate how *F* affects the dynamical properties of chromatin. The movements of the loci in the gene-rich region are suppressed in an intermediate range of *F*, and are enhanced at small *F* values, which has also been observed in experiments. In the intermediate *F*, the bond length between consecutive loci increases, becoming commensurate with the distance at the minimum of the attractive interaction between non-bonded loci. This results in a transient disorder-to-order transition, leading to the decreased mobility during transcription. Strikingly, the *F* -dependent change in the locus dynamics preserves the organization of the chromosome at *F* = 0. Transient ordering of the loci, which is not found in the polymers with random epigenetic profiles, in the gene-rich region might be a plausible mechanism for nucleating a dynamic network involving transcription factors, RNAPII, and chromatin.

**Significance Statement:** In order to explain a physically counter-intuitive experimental finding that chromatin mobility is reduced during transcription, we introduced a polymer model for interphase chromosome that accounts for RNA polymerase (RNAP) induced active force. The simulations show that, over a range of active forces, the mobility of the gene-rich loci is suppressed. Outside this range, chromosomes are compact and exhibit glass-like dynamics. Our study, which accounts for the experimental observations, leads to a novel and testable mechanism of how transcription could shape the coexistence of fluid- and solid-like properties within chromosomes.

---

Advances in experimental techniques (1, 2) have elucidated the organizational details of chromosomes, thus deepening our understandings of how gene regulation is connected to chromatin structure (3). Relatively less is known about the dynamics of the densely packed interphase chromosomes in the cell nucleus. Experimental and theoretical studies have shown that the locus dynamics is heterogeneous, exhibiting sub-diffusive behavior (4–8), which is consistent with models that predict glass-like dynamics (9–11). However, it is challenging to understand the dynamic nature of chromosomes that governs the complex subnuclear processes, such as gene transcription.

The link between transcriptional activity and changes in chromosome dynamics is important in understanding the dynamics of chromosomes in distinct cell types and states (12, 13). It is reasonable to expect that transcription of an active gene-rich region could make it more expanded and dynamic (14, 15). However, active RNA polymerase (RNAP) II suppressed the movement of individual loci in chromatin (13, 16). Let us first summarize the key experimental results (13, 16), which inspired our study: (1) By imaging the motion of individual nucleosomes (monomers in the chromosomes) in live human cells, it was shown that the mean square displacements (MSDs) of the nucleosomes during active transcription are constrained (see Fig. 1A). (2) When the cells were treated with *α*-amanitin (*α*-AM) or 5,6-Dichloro-1-*β*-D-ribofuranosylbenzimidazole (DRB), transcription inhibitors which selectively block the translocation of RNAPII (17, 18), the mobilities of the nucleosomes were enhanced (Figs. 1A– 1B). This finding is counter-intuitive because the translocation generates mechanical forces (19, 20) that should render the chromatin more dynamic. (3) The enhanced motion was restricted only to gene-rich loci (euchromatin) that are predominantly localized in the cell interior (21, 22) whereas the dynamics of gene-poor heterochromatin, found in the periphery (21–23), is unaffected by transcription. Based on these observations, it was hypothesized that RNAPs and other protein complexes, facilitating transcription, transiently stabilize chromatin by forming dynamic clusters (24–28), whose structural characteristics are unknown. This hypothesis has been challenged because the inhibition mainly leads to stalling of RNAPs bound to chromatin (17, 18). Moreover, transcriptional inhibition does not significantly alter the higher-order structures of chromosomes (28, 29). These observations raise the question: Is there a physical explanation for the increased chromatin dynamics upon inhibition of transcription and a decrease during transcription? We provide a plausible answer to this question by using extensive simulations of a simple minimal active copolymer model, which captures the organization of chromosomes in the absence of active forces.

**Fig. 1.**
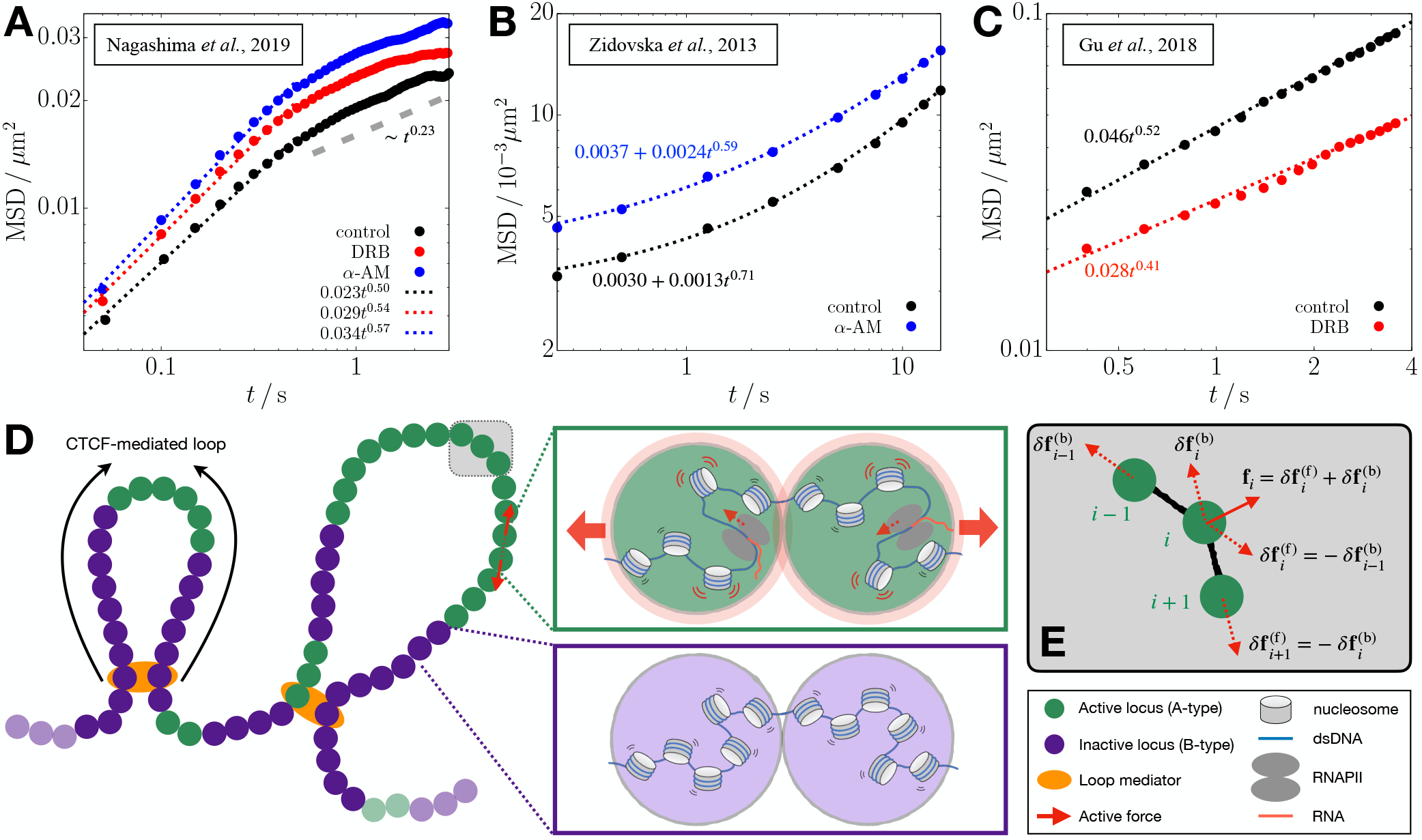
Comparison of chromatin mobility changes in different experiments of transcription inhibition (top) and the schematic depiction of the polymer model used in the current study (bottom). (A–C) Time-dependent MSD data from the experiments of Refs. 13 (A), 16 (B), and 14 (C) are plotted on a log scale. The data for the nuclei treated with DRB and *α*-amanitin (*α*-AM) are shown in red and blue circles, respectively, whereas those for the control (untreated nuclei) are in black circles. The dotted lines are the best fits to the data in the given time range (*t <* 0.5 s for panel A). The gray dashed line in panel A shows the scaling of MSD for *t >* 0.5 s suggestive of the constrained motion of nucleosomes. (D) Chromosome is modeled as a copolymer chain consisting of active/inactive (green/purple-colored) loci. Specific locus pairs are connected by loop mediators(orange color). We apply active forces to all the gene-rich A-type loci in the form of an extensile force dipole (red arrows) along each bond vector between the A-type loci.The green and purple boxes illustrate the microscopic origin of the dipolar extensile active forces. Compared to the inactive B-type loci (bottom), the A-type loci include the nucleosomes in a more dynamic state along with the actively transcribing RNAPII, such that their effective excluded volumes increase modestly, as represented by the light-red shade. The enhanced volume exclusion gives rise to repulsion between the consecutive A-type loci and the increase of the bond distance. (E) Illustration for how active forcesare imposed on the *i*th locus and its bonded loci, where the solid arrow is the net force from the force dipoles (dashed arrows).

In contrast to the experimental findings discussed above, it has been shown that under certain conditions (14) the mobility of gene loci increases in the presence of transcription. In other words, the mobility decreases when transcription is inhibited (see Fig. 1C). This intuitive result may be ratinoalized by assuming that transcription exerts active forces although differences between the results in Figs. 1C and 1A are hard to explain.

Using the experimental results as a backdrop, we theorize that RNAPII exerts active force (19, 20) in a vectorial manner on the active loci. We then examine the effects of active force on the organization and dynamics of chromosome using the Chromosome Copolymer Model (CCM) (10). In the absence of active forces, the CCM faithfully captures the experimental results, showing microphase separation between euchromatin (A-type loci) and heterochromatin (B-type loci) on large length scale and formation of substructures on a smaller length scale in interphase chromosomes. The CCM leads to a condensed polymer, exhibiting glass-like dynamics in the absence of activity, in agreement with experiments (4, 5, 10, 30). Brownian dynamics simulations of a single chromosome in the presence of active forces, produces the following key results: (i) In accord with experiments (13), the dynamics of the active loci, measured using the MSD, is suppressed upon application of the active force. Despite the use of minimal model, our simulations qualitatively capture the increase in the MSD for the transcriptionally inactive case relative to the active case. (ii) The decrease in the mobility of the A loci occurs only over a finite range of activity level. Surprisingly, in this range the segregated A loci undergo a transient disorder-to-order transition whereas the B loci remain fluid-like. (iii) Chromosome in which the epigenetic (A/B) profile is random cannot capture the experimental observations, implying that the sequence inherent in the chromosomes plays a vital role.

## Model

### Chromosome Copolymer Model

We model an interphase chromosome as a flexible self-avoiding copolymer (Fig. 1D), whose potential energy function is given in Methods. Non-adjacent pairs of loci are subject to favorable interactions, modeled by the Lennard-Jones (LJ) potential (Eq. 6), depending on the locus type. The relative strengths of interactions between the loci is constrained by the Flory-Huggins theory (31, 32), which ensures microphase separation between the A and B loci—an important organization principle of interphase chromosomes. Certain locus pairs, separated by varying genomic distances, are linked to each other, representing the chromatin loops mediated by CCCTC-binding factors (CTCFs) (33). In this study, we simulate a 4.8-Mb segment of human chromosome 5 (Chr5: 145.87–150.67 Mb) using *N* = 4,000 loci, that is,1.2 kb *∼* 6 nucleosomes per locus. The A-to B-type ratio is *N*_A_*/N*_B_ = 982*/*3018 *≈* 1*/*3. See Methods for the details about assignment of the locus type and the loop anchors in the CCM polymer.

In our model, the loop topology does not change with time, unlike polymer models, which examined the consequences of dynamic extrusion of loops on interphase and mitotic structures (34–36). We simulate a given chromosome region up to *∼*10 seconds (see Methods for details) that is much shorter than the lifetime of the CTCF loops (15–30 mins) (37, 38). Furthermore, the loop extrusion rate is *∼* 0.5–2 kb/s (39, 40), which also would not significantly alter the topology of loops on the simulation time scales. Therefore, the assumption that loops are static is reasonable. We discuss, in a later section, the potential effects of the removal of loops along with the implications (see Discussion)

### Active forces in transcriptional activity

Previous theoretical studies (41–49) have considered different forms of active forces on homopolymers and melts of polymer rings. Potential connection to chromatin has been proposed using polymer models in the presence of active forces (6, 50–55). However, these have not accounted for the counter-intuitive finding from the experiment of transcription inhibition (13). Here we applied active forces on the CCM polymer chain to mimic the force generated by the actively transcribing RNAPII during transcription elongation (19, 20).

In our model, active forces act along each bond vector between the A-type loci in an extensile manner, ensuring momentum conservation (see Fig. 1D). The force, associated with the translocation of actively transcribing RNAPII (19, 20), should result in the increased volume exclusion between the bonded A loci, as shown in the green box in Fig. 1D. We reason that the dipolar extensile active force, with local momentum conservation, models the effective repulsion between the active loci in the coarse-grained level (see SI Appendix, Sec. 1 for more details about the biological relevance of the force). Along the A-A bond vector, **b**_*i*_ = **r**_*i*+1_ *−* **r**_*i*_ (**r**_*i*_ is the position of the *i*^th^ locus), force, 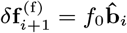, acts on the (*i* + 1)^th^ locus in the forward direction (*f*_0_ is the force magnitude and 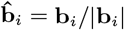, and 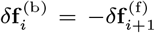 is exerted on the *i*^th^ locus in the opposite direction (Fig. 1E). If the (*i−*1)^th^ locus is also A-type, another pair of active force is exerted along the bond vector, **b**_*i −* 1_ = **r**_*i*_ *−***r**_*i −*1_, and thus the net active force on the *i*^th^ locus is given by 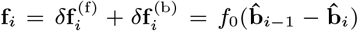 (solid arrow in Fig. 1E). We simulate the CCM chain dynamics by numerically solving the overdamped Langevin equation, given in Eq. 10, which includes the active forces (see Methods for simulation details). We use the dimensionless parameter, *F≡f*_0_*σ/k*_*B*_ *T*, as a measure of the force magnitude, where *σ* is the diameter of a single locus, *k*_*B*_ is the Boltzmann constant, and *T* is the temperature.

In the simulations, we apply the active forces of given magnitude *F* to all the A-type loci at each time step. Our choice of the locus resolution (1.2 kb) is sufficiently small to distinguish between the gene and the intergenic loci which are assigned A- and B-types, respectively (note that the average human gene size is*∼*27 kb (56)). Here we assume that all the genes in the simulated chromosome region are co-activated, and experience the same magnitude of forces during our simulation. In a later section, we discuss the potential effects of changing the gene activations and the corresponding force exertion (see”Connection to other experiments” in Discussion).

On each gene body, there are multiple active RNAPII complexes during a transcriptional burst, the time in which transcripts are generated (57, 58). For a human gene, the number of RNAPII or transcribed RNAs is observed to be up to 5 during the burst interval of 10–20 minutes, which seems to be conserved across other genes (59). Hence, the average linear density of active RNAPII in each gene during the burst can be estimated as*∼* 0.01–0.05 RNAPII per kb. On the other hand, RNAPII clusters observed in live-cell imaging experiments (25, 60) have the volume density of *∼*10^4^ RNAPII per *μ*m^3^, which corresponds to*∼*1 RNAPII per kb based upon genome density in the nucleus. Although the clusters are closely related to transcriptional activity (25, 61), it is unclear what fraction of RNAPII in the clusters is activated and engaged in the transcription elongation of genes. Moreover, it is unknown what kind of forces are generated on mammalian chromatin during transcription, which is crowded with RNAPII and other protein factors. Here we explore the dynamics of the model chromatin subject to varying active forces.

In the following, we first present the simulation results where the active forces were applied on all the A-type loci. In the Discussion the section, we compare the main results with those from the simulations with lower force density (see “Effect of active force density”).

## Results

### MSD comparison

We calculated the MSDs separately for euchromatin and heterochromatin loci at lag time *t*,

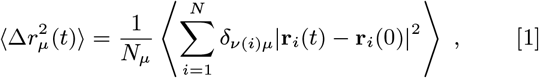

where ⟨ *· · ·* ⟩ is the ensemble average, *ν*(*i*) is the type of the *i*^th^ locus, and *δ*_*ν*(*i*)*μ*_ is the Kronecker delta (*δ*_*xy*_ = 1 if *x* = *y*, or 0 else) that picks out the loci of specific type *μ* (either A or B). Fig. 2A shows that 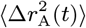 is smaller at *F* = 80 compared to *F* = 0. This result is qualitatively similar to the nucleosome MSDs measured from the interior section of the cell nucleus treated with the transcription inhibitor *α*-AM (13). 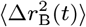 at *F* = 80 is also smaller than at *F* = 0, but the difference is marginal compared to 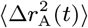, which is consistent with the experimental results for the nuclear periphery (Fig. S1A). In Fig. 2A, the magnitude of 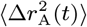 is smaller in simulations than in experiment, where the difference in magnitude depends on the conversion of simulation length/time to real units (see Methods for the details). The level of agreement with experiments is striking, considering that we used a minimal model with no parameters to describe a very complicated phenomenon. We could probably have obtained better agreement with experiments by tweaking the parameters in the model, which we think is not necessary for the purposes of this study.

**Fig. 2.**
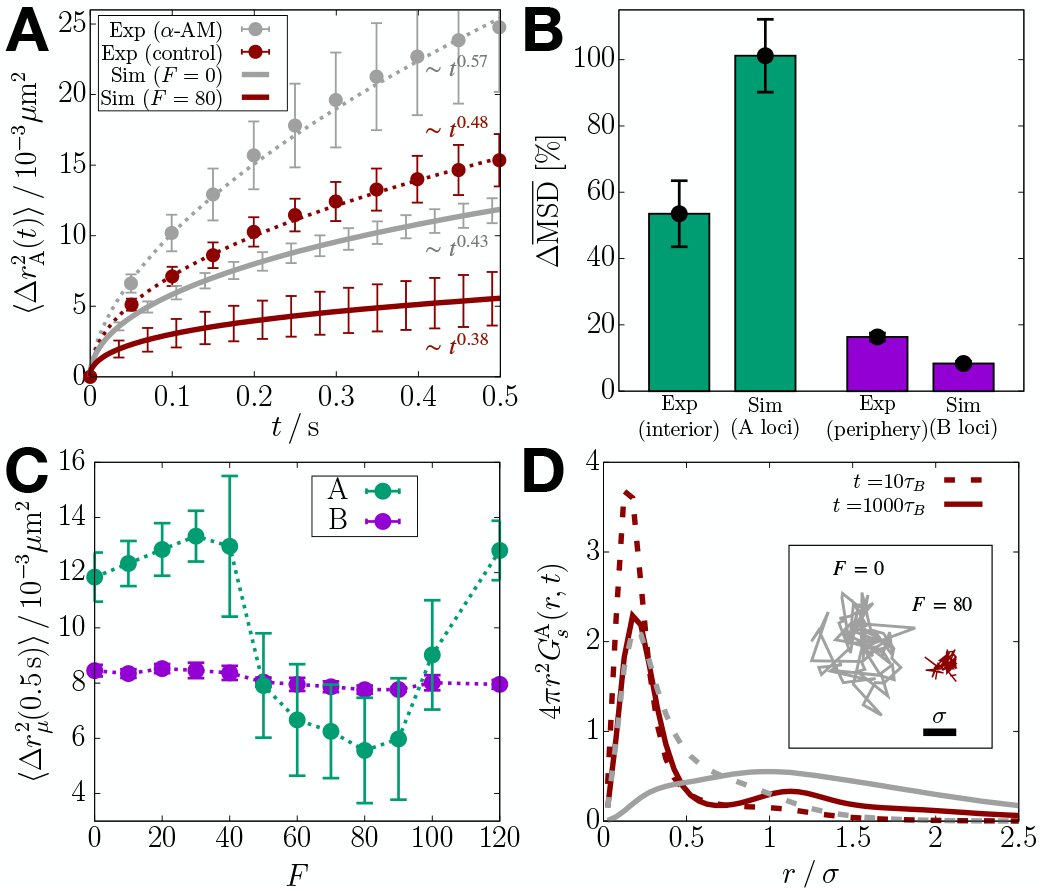
Transcription-induced active forces reduce the mobilities of euchromatin loci. (A) 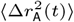 from simulations with F = 0 and F = 80 (solid lines), compared with the euchromatin MSD from the experiment that inhibits transcription using α-AM (13) (circles). The dotted lines are the fits to the experimental data. The error bars for the simulation data are obtained from five independent trajectories. (B) Bar graphs comparing the increase in 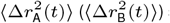 shown in panel A (Fig. S1A) between experiment and simulation results. (C) MSDs for the A and B loci at t = 0.5 s as a function of F. The dotted lines are a guide to the eye. (D) Radial distributions of the A-locus displacement at t = 10_τB_ (dashed) and t = 1,000_τB_ (solid), compared between F = 0 (gray) and F = 80 (dark-red). The inset shows the 2-D projectionof the trajectory of an active locus for 10^4^_τB_ at F = 0 and F = 80.

In Fig. 2A, each MSD plot as a function of time is shown with scaling behavior,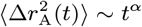. The scaling exponent, *α*, provides the link between chromatin structure and the mobility. It was established previously (6) that, 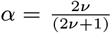 where *ν*, the Flory exponent related to the radius of gyration(*R*_*g*_ ∼*N* ^*ν*^)of the polymer chain. For an ideal Rouse polymer, *ν* = 0.5, and for a polymer in a good solvent, which takes the excluded volume between the monomers into account, *ν ≈* 0.6 (62). The fit to the experimental data (gray dotted line in Fig. 2A) when transcription is inhibited yields *α ≈* 0.57, which would be consistent with the prediction for the SAW behavior. In all other cases, *α* is less than 0.5 (the value for the Rouse polymer) both in experiments and simulations, which is indicative of restricted motility. The scaling exponent in the experimental MSD increases from *α* = 0.48 (*±*0.05) to *α* = 0.57 (*±* 0.07) upon inhibiting transcription. We estimated the errors using 10,000 samples reconstructed from the normal distribution for each data point, following the procedure described elsewhere (63). Our simulations capture the change in *α* such that Δ*α*(active *→* passive) = 0.05 (versus 0.09 from the experiments). In contrast, recent experiments for chromatin dynamics in different contexts, such as loop formation and response to mechanical perturbation, reported that the MSD exponents are similar to the Rouse (37, 64) or SAW (38) polymer, *i*.*e*., *α* ≳ 0.5. These results imply fundamentally different physical nature in transcriptionally active regions, which constrains the polymer dynamics to *α <* 0.5.

In Fig. 2B, we compare the transcription-inhibited increase 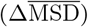 in the MSD, between experiment and simulations (see Eqs. S1–S2 in SI Appendix). We use *F* = 80, which shows the smallest MSD (see Fig. 2C), as the control. The value of *f*_0_ for *F* = 80 is in the range, *f*_0_*≈* 3–16 pN (see Methods), which accords well with the typical forces exerted by RNAP (19). Comparison between 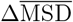 for the A loci (simulation) and the interior measurements (experiment) is less precise than between the B loci and the periphery. The difference may arise because the interior measurements could include the heterochromatin contribution to some extent, whereas the periphery measurements exclude the euchromatin. Nevertheless, 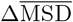 for all the loci is in better agreement with experiment (Fig. S1C). The reasonable agreement between simulations and experiment is surprising because it is obtained *without adjusting any parameter to fit the data*. Although comparisons in Fig. 2B are made with *F* = 80, we obtain qualitatively similar results for *≤F* in the range, *≤* 60 *F* 90 (Fig. S2).

The simulated MSD, at a given lag time, changes non-monotonically as *F* changes. Remarkably, the change is confined to the A loci (Figs. 2C and S1D–S1E); 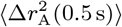 increases modestly as *F* increases from zero to *F* ≲ 30, and decreases when *F* exceeds thirty. There is an abrupt reduction at *F ≈* 50. In the range, 50 ≲ *F* ≲ 80, 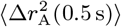 (0.5 s) continues to decrease before an increase at higher *F* values. In contrast, the MSD of the B loci does not change significantly with *F* (Fig. 2C). The non-monotonic trend in 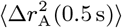 upon changing *F* is reminiscent of a *re-entrant* phase behavior in which a system transitions from an initial phase to a different phase and back to the initial phase as a parameter (*F* in our case) is varied. Such a behavior occurs in a broad range of different systems (65–68). We present additional analyses in the next section to confirm the *dynamic re-entrance*.

To obtain microscopic insights into the simulated MSDs, we calculated the van Hove function, which is simply the distribution of locus displacement at a given lag time,

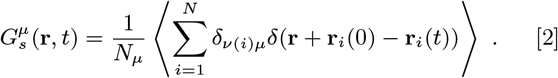

Figure 2D compares the radial distribution for the displacement of A-type loci, 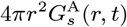, between *F* = 0 and *F* = 80 at *t* = 10*τ*_*B*_ *≈* 0.007 s and 1,000*τ*_*B*_ 0.*≈* 7 s (see Methods for the unit conversion). At short lag times, both the distributions for *F* = 0 and *F* = 80 show a peak at *r <* 0.5*σ*, which means that a given locus is unlikely to diffuse far from the neighboring loci in a condensed microenvironment. At long lag times, the distribution for *F* = 0 becomes more populated at *r≥σ*, as the loci escape from the regime caged by the neighbors. The remarkably broad and flat curve for 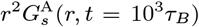 at *F* = 0, resembling a uniform distribution, signifies the heterogeneous dynamics of the A-type loci, which differs from the Gaussian distribution expected for the Rouse polymer. The A-type loci at *F* = 80 do not diffuse as much at *F* = 0 even at long lag times (Fig. 2D), and their displacements are largely within the length scale of *σ*. In contrast, there is no significant difference in 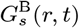 between *F* = 0 and *F* = 80 (Fig. S1F). Notably, the second peak of 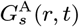 at *F* = 80 hints at the solid-like lattice (69).

### Dynamic re-entrant behavior

Here, we further characterize the dynamics of the CCM polymer chain in the presence of RNAPII-induced active forces. The purpose is to investigate how the re-entrant behavior emerges in the CCM polymer whose dynamics is glass-like in the absence of the activity (10). To probe how the glass-like behavior of the polymer chain (6, 9, 10) is affected by the active forces, we calculated the self-intermediate scattering function,

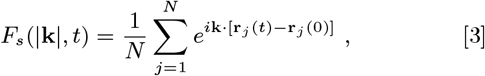

where **k** is the wave vector. The scattering function, the Fourier transform of the van Hove function (Eq. 2), quantifies the extent of the loci displacements over time *t* relative to the length scale set by the wave vector (*∼*2*π/*|**k**|). We computed the ensemble-averaged ⟨ *F*_*s*_(*k*_max_, *t*)⟩ with 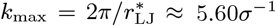, where 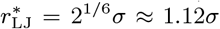 is the distance at the minimum of the non-bonding (LJ) potential (see Methods),which the nearest neighbors are likely to be located at (see the radial distribution function in Fig. 4 below). The decay of ⟨*F*_*s*_(*k*_max_, *t*) ⟩ indicates the structural relaxation on the length scale ≳ *σ*.

Time-dependent variations in ⟨*F*_*s*_(*k*_max_, *t*)⟩ (Fig. 3A) show stretched exponential behavior 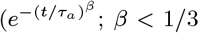 at all *F* values), which is one signature of glass-like dynamics. Note that the decay is even slower if *F* is increased, which is in agreement with the results shown in Figs. 2 and S1. The relaxation time, *τ*_*α*_, calculated using ⟨*F*_*s*_(*k*_max_, *τ*_*α*_)⟩ = 0.2, shows that the relaxation is slowest at *F ≈* 80 (Fig. 3B), which occurs after the dynamical transition in 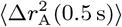 at *F* ≈ 50 and before 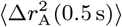 increases beyond *F* = 100 (Fig. 2C). Similarly, when the tails of ⟨*F*_*s*_(*k*_max_, *t*)⟩ were fit with 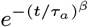, the exponent *β* also exhibits the analogous trend (Fig. 3B); that is, as *τ*_*α*_ increases, *β* decreases, indicating the enhancement in the extent of the glass-like behavior.

**Fig. 3.**
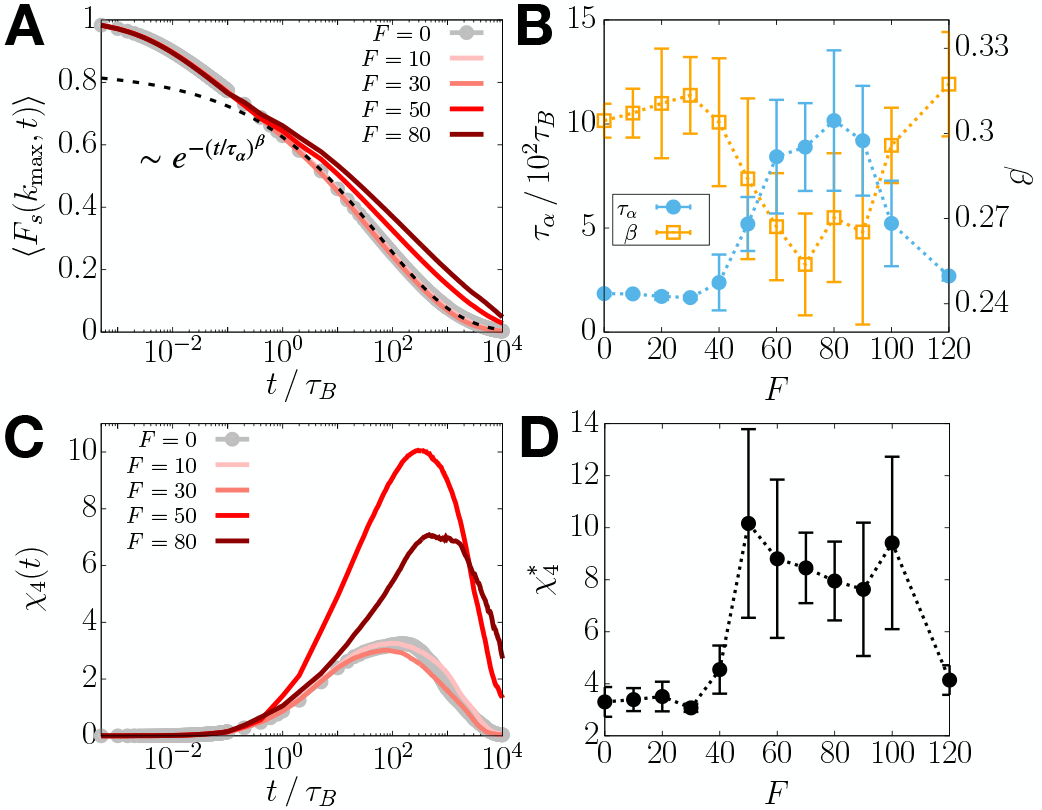
Re-entrance in the relaxation time and dynamic heterogeneity is observed upon increasing the activity. (A) Plot of ⟨F_s_(k_max_, t)⟩ (Eq. 3) for different F. The dashed line is a stretched exponential fit for F = 0. (B) τα (blue) and β (orange) of ⟨F_s_(k_*max*_, t)⟩ as a function of F. The dotted lines are a guide to the eye. (C) Plot of χ_4_(t) (Eq. 4) for different F. (D) The maximum value of χ_4_(t) as a function of F.

**Fig. 4.**
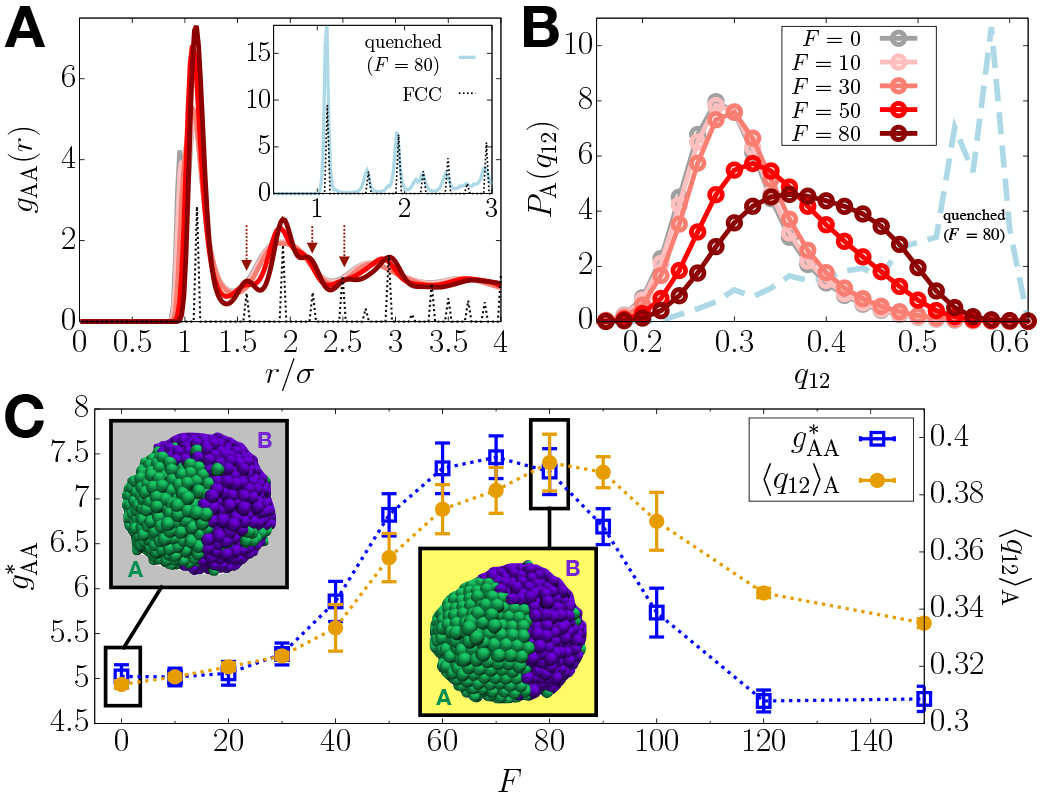
*F* -induced structural transition. (A) RDF for A-A pairs at different *F* (solid lines; see the legend in panel B), where *g*(*r*) for a FCC crystal is shown with the dotted line (scaled arbitrarily). The inset shows *g*_AA_(*r*) for the quenched polymer with *F* = 80. (B) Distributions of the BOO parameter, *q*_12_, for A loci as a function of *F*. The dashed line is for the quenched A loci at *F* = 80. (C) Height of the dominant peak in *g*_AA_(*r*) (blue) and ⟨*q*_12_⟩ _A_ (orange) as a function of *F*. Simulation snapshots for *F* = 0 and *F* = 80 are in the gray and yellow boxes, respectively.

Dynamic heterogeneity, another hallmark of glass-like dynamics (70, 71), was calculated using the fourth-order susceptibility (72), which is defined as the mean squared fluctuations in the scattering function,

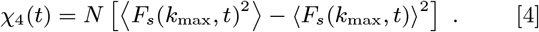

We find that *χ*_4_(*t*), at all *F* values, has a broad peak spanning a wide range of times, reflecting the heterogeneous motion of the loci (Fig. 3C). The peak height, 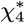, increases till *F ≈* 50 and subsequently decreases (Fig. 3D). When *F* exceeds 100, 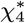 decreases precipitously. Our results suggest that there are two transitions: one at *F ≈* 50 where the dynamics slows down and the other, which is a reentrant transition beyond *F* = 100, signaled by an enhancement in the mobilities of the A-type loci. Although the system is finite, these transitions are discernible.

Like the MSD, when ⟨*F*_*s*_(*k*_*max*_, *t*) ⟩ and *χ*_4_(*t*) was decomposed into the contributions from A and B loci (Eqs. S3 and S4), we find that the decrease in the dynamics and the enhanced heterogeneity are driven by the active loci (Fig. S3). These observations, including the non-monotonicity in *τ*_*α*_ and *β* that exhibit a dynamic reentrant behavior, prompted us to examine if the dynamical changes in the A-type loci are accompanied by any structural alterations.

### Transient disorder-to-order transition induced by active forces

The radial distribution function (RDF) for A-A locus pairs, *g*_AA_(*r*), with signature of a dense fluid, shows no visible change for *F* ≲ 30 (Fig. 4A). In contrast, the height of the primary peak, 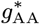, increases sharply beyond *F* = 30 (Fig. 4C). Remarkably, *g*_AA_(*r*) for *F* = 80 exhibits secondary peaks that are characteristics of the FCC (face-centered cubic) crystal phase (Fig. 4A, arrows). Upon further increase in *F*, these peaks disappear (Fig. S4A) and 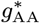 reverts to the level of the passive case (Fig. 4C). In other words, the active forces, which preserve fluid-like disorder in the A loci at low *F* values, induce a structural transition to solid-like order in the intermediate range of *F* values, which is followed by reentrance to a fluid-like behavior at higher *F* values. In contrast, *g*_BB_(*r*) exhibits dense fluid-like behavior at all *F* values (Fig. S4B). We confirm that the FCC lattice is the minimum energy configuration by determining the *inherent structure* (73, 74) for the A loci at *F* = 80 by quenching the active polymer to a low temperature (see Methods) (Fig. 4A, inset). In Fig. S4C, *g*_AA_(*r*) at *F* = 0 reflects the structure of a dense fluid upon the quench whereas *g*_AA_(*r*) at *F* = 80 is quantitatively close to that for a perfect FCC crystal. Quenching does not alter the structure of the B loci at *F* = 80 or *g*_AA_(*r*) at *F* = 0 (Figs. S4C-S4D).

To assess local order, we calculated the bond-orientational order (BOO) parameter for 12-fold rotational symmetry, *q*_12_ (Eqs. 12–13) (75, 76). For a perfect FCC crystal, *q*_12_ *≈* 0.6 (77). The distribution for A loci, *P*_A_(*q*_12_) (Eq. 14), is centered around *q*_12_ = 0.3 at *F* = 0 (Fig. 4B), representing a disordered liquid state (Fig. 4C, gray box). As *F* increases, the distribution shifts towards the right, especially in the 50 *≤ F ≤* 80 range. The increase of ⟨*q*_12_ ⟩_A_ (Eq. 15) indicates a transition to a FCC-like ordered state that is visible in the simulations (Fig. 4C, yellow box). *P*_A_(*q*_12_) at *F* = 80 is broad, whereas the inherent structure gives a narrower distribution peaked near *q*_12_ = 0.6 (Fig. 4B, dashed line), which shows that the ordered solid-like state coexists with the disordered fluid-like state within thermal fluctuations. The maximum in *P*_A_(*q*_12_) shifts to the left for *F >* 80 (Fig. S4E) and ⟨*q*_12_⟩_A_ decreases, suggestive of *F* -induced reentrant transition. The distribution *P*_B_(*q*_12_) for the B-type loci is independent of *F* (Fig. S4F). These results show that FCC-like ordering emerges in 50 ≲ *F* ≲ 100 range. Outside this range, the RDFs display the characteristics of a dense fluid. In addition, the *F* -dependent bending angle distributions for three consecutive A loci (Eq. S5) reflect the FCC-like ordering (Fig. S5). The transitions in the A-type loci may be summarized as fluid *→* FCC/fluid *→* fluid, as *F* changes from 0 to 120.

Since the overall RDF at *F* = 80 does not show the FCC peaks (Fig. S6A), the *F* -induced order may not be seen directly in the conventional experiments. Moreover, the ordering is transient, as implied in the coexistence of order and disorder. By transient we mean that solid-like order persists only when the polymerase induces active forces during transcription. The time correlation function of the BOO parameter,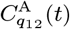 (Eq. S6), decays rapidly, at most on the order, *≈* 0.7 s (Fig. S6B), much shorter than the transcription time scale (*∼* minute). More details and the visualization of the transient ordering are given in SI Appendix, Sec. 5 and Movie S1. Remarkably, the transient ordering preserves the large-scale structure of chromosome (see SI Appendix, Sec. 6; Fig. S7), in agreement with the chromosome conformation capture experiments with transcription inhibition (28, 29). Here we emphasize that the solid-like nature coexisting with the liquid is sufficient to explain the experimentally observed suppression of euchromatin motions. The predicted transient ordering could be detected by future high resolution imaging experiments.

### Origin of *F* -induced order

The emergence of solid-like order in the A-type loci is explained using the effective A-A interaction generated by *F*. We calculated the *effective pair potential* for an A-A bond,

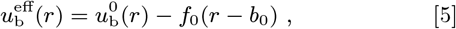

where 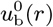 and *b*_0_ are the *F* -independent bonding potential, and the corresponding equilibrium bond length, respectively. The *f*_0_(*r* − *b*_0_) term is the work done by the active force to stretch the bond from *b*_0_. The equation of motion for *F* ≠ 0 represents dynamics in the *effective equilibrium* under the potential energy involving 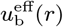 (Eqs. 16–17). Such effective equilibrium concept was useful for characterizing various physical quantities in active systems (47, 54, 78–81).

Plots of 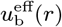 in Fig. 5A show that the effective equilibrium bond length, *r*_min_, increases as *F* increases. This prediction is confirmed by the direct measurement of A-A bond distance from the simulations (Figs. S8A–S8B). Note that 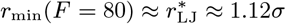, where 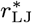 is the distance at the minimum of the LJ potential (Fig. 5B). The *F* -induced extension of A-A bonds makes *r*_min_ commensurate with 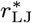, which is conducive to FCC ordering (82) in the active loci. At *F* = 0, since the bond distance (*b*_0_ = 0.96*σ*; see Methods) is smaller than the preferred non-bonding interaction distance, 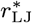, the lattice-matching configurations cannot be formed given the chain connectivity (Fig. 5C, left). In contrast, at *F* = 80, the condition that 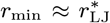 allows the A loci to be arranged into the FCC lattice, leading to the transition to the ordered state (Fig. 5C, center). Upon increasing the activity further (*e*.*g*., *F* = 120), the bond distance becomes too large to be fit with the lattice structure, so the loci re-enter the disordered fluid state (Fig. 5C, right).

**Fig. 5.**
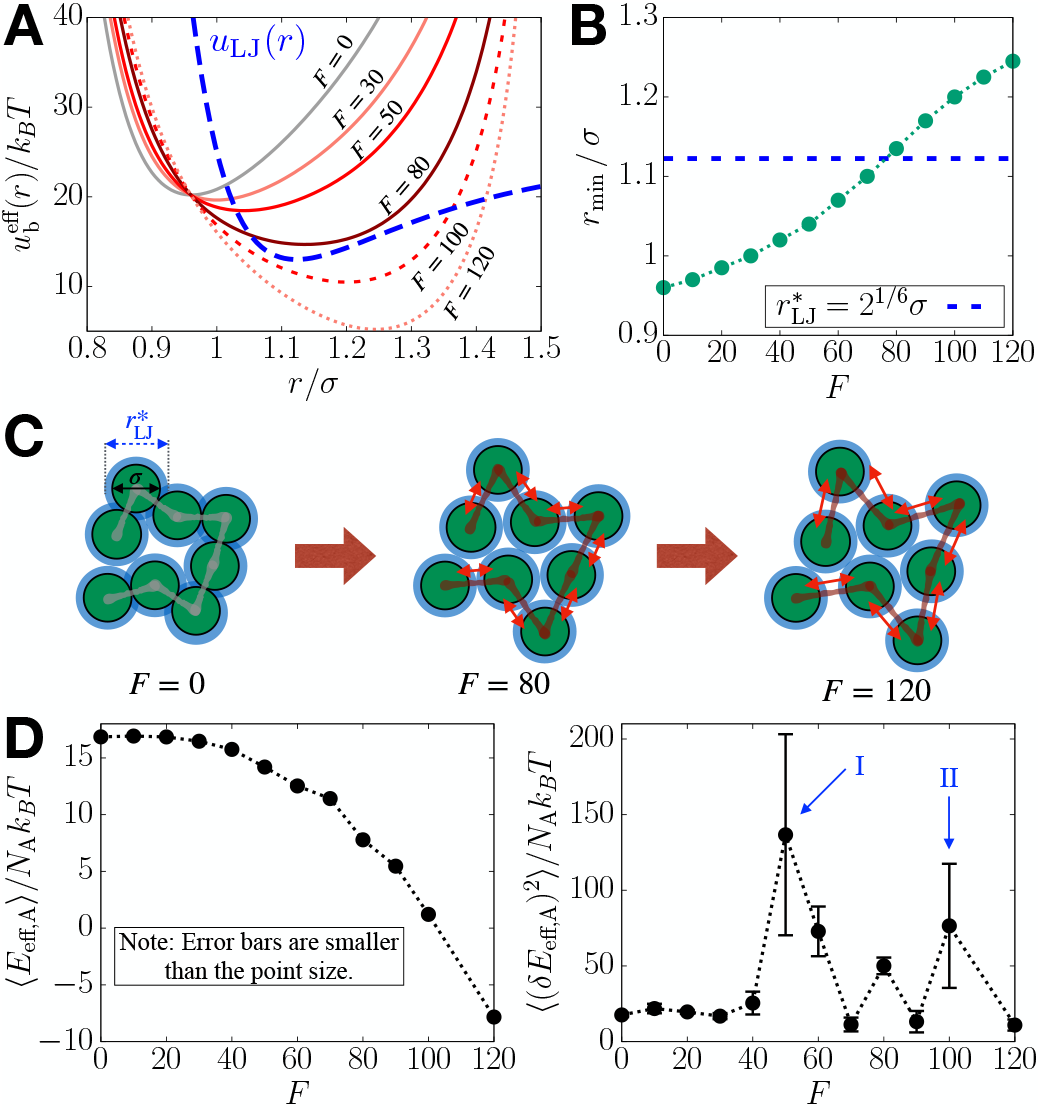
Effective potential energy accounts for the structural transition of the A-type loci. (A) Effective pair potential of a single A-A bond, 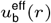 (Eq. 5), as a function of *F*. The LJ potential, *u*_LJ_(*r*), which is vertically shifted, is shown in blue dashed line for comparison. (B) Distance at the minimum of 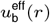 as a function of *F*, where the dashed line indicates 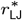 (see the main text). (C) Schematic illustration for how the *F* - induced bond extension enables FCC ordering in the A-type loci (green circles). The blue shade shows 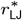 at which non-bonded pairs have the most favorable interactions. The double red arrows indicate the bond extension upon increasing *F*. (D) Mean of the effective potential energy of A loci (left) and mean fluctuations (right) with respect to *F*. The arrows indicate the two structural transitions.

We can also describe the ordering behavior using thermodynamic properties based on 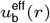. We calculated the mean and variance of the effective potential energy of the A loci, ⟨*E*_eff,A_⟩ (Eq. 18). Fig. 5D shows that *E*_eff,A_ decreases smoothly as *F* changes, without pronounced change in the slope, as might be expected for a structural transition (82). Nevertheless, ⟨ (*δE*_eff,A_)^2^⟩ indicates signatures of a transition more dramatically, with peaks at *F* = 50 and *F* = 100 (Fig. 5D, arrows I and II). Thus, both ordering and reentrant fluid behavior coincide with the boundaries of the dynamic transitions noted in Figs. 2B, 3B, and 3D.

## Discussion

We introduced a minimal active copolymer model for Chr5 in order to explain the non-intuitive experimental observation that during transcription the mobilities of the loci are suppressed. Despite the simplicity of the polymer model, we reproduce semi-quantitatively the experimental observations. In particular, the MSD exponent 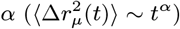 decreases by 0.05 in the simulations whereas the decrease is 0.09 between the transcription inhibited and active states in the experiments (Fig. 2A). We find this result and the results in Fig. 2 surprising because no parameter in the very simple model was adjusted to obtain the semi-quantitative agreement with experiments (13).

We hasten to add that the model indeed is oversimplified, and is the major limitation of our study. The actual machinery of transcription active (or inhibited) state is extremely complicated involving the interplay of transcription factors, RNAPII, chromatin, and other cofactors. These elements are believed to drive the formation of a hub of connected modules resulting in condensates (24–27). Although the structural features of chromatin in the condensates are unknown, our simulations suggest that for a very short period (*≈* 1s) chromatin could adopt solid-like properties when RNAPII and other transcription-associated machinery exert an active force (*≈* 5–10 pN) during transcription.

### Robustness of the conclusions

We performed three tests in order to ensure that the results are robust. (1) Simulations of a segment of chromosome 10, with a larger fraction of active loci, show qualitatively similar behavior (See SI Appendix, Sec. 7; Fig. S9). (2) For a copolymer chain whose A/B sequence is random (See SI Appendix, Sec. 8; Fig. S10), we find that at all *F*, MSD for the A-type loci increases monotonically (Fig. 6A) in contrast to the non-monotonic behavior found in Chr5 with a given epigenetic (A/B) profile (Fig. 2C). The B-type MSD decreases at high *F* (Fig. 6A), which is also different from the behavior in Chr5. In addition, in the random sequences the loci exhibit fluid-like behavior at all values of *F*, and hence cannot account for the experimental observation (Fig. 6B). Thus, *F* -induced decrease in the motility of the A loci, accompanied by transient ordering, occurs only in copolymer chains exhibiting microphase separation between A and B loci—an intrinsic property of interphase chromosomes (83). In other words, the epigenetic profile determines the motilities of the individual loci (or nucleosomes) during transcription. (3) We performed simulations by removing all the loop anchors (analog of topological constraints (47); see SI Appendix, Sec. 9). Fig. S11 shows that the *F* -induced dynamical slowdown and ordering is not significantly affected by the loop anchors. The modest increase in the MSD upon removal of loops, at *F* = 0, is qualitatively consistent with the experimental measurements of increased chromatin mobility when cohesin loading factors were deleted (12). The experimental results suggest that a reduction in transcriptional activity may be induced by loss of loops, which is assumed to be a consequence of depletion of cohesin (see “Connection to other experiments” below for more discussion). These tests show that our conclusions are robust, and the model might serve as a minimal description of the effect of transcription-induced active forces on real chromosomes.

**Fig. 6.**
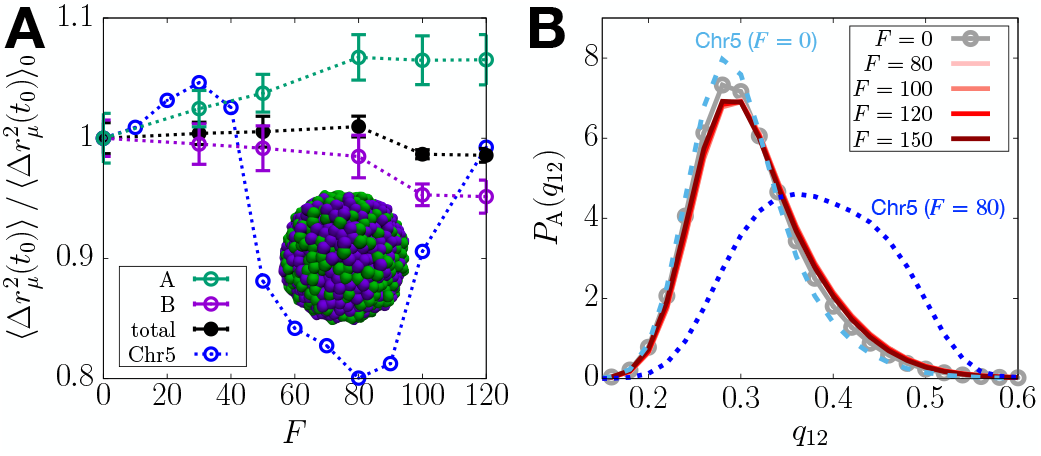
A copolymer chain with the same length and the fraction of active loci as Chr5 but randomly shuffled epigenetic profile does not capture the experimental observations. (A) Ratio of the MSD at *t*_0_ = 100*τ*_*B*_ for the active random copolymer to the passive case as a function of *F*, shown in black color. The data for Chr5 is shown in blue color. The same quantities computed for the A and B loci of the random copolymer are also plotted in green and purple color, respectively. The dotted lines are a guide to the eye. The simulation snapshot shows that the copolymer chain is in a mixed state without discernible microphase separation. (B) Probability distributions of *q*_12_ for the A loci at different activity levels, where the dashed and dotted lines are the distributions for Chr5 with *F* = 0 and *F* = 80, respectively.

### Emergence of solid-like ordering induced by active forces

In the single copolymer chain, which reflects the compact nature of interphase chromosomes (1, 84), transient solid-like ordering would occur only if the distance between two bonded loci is roughly equal to the distance at which the pair potential between the non-bonded loci is a minimum (for the LJ potential,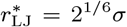) (82). This argument is generic because it only depends on the presence of attractive interactions between non-bonded loci, which is needed to ensure that interphase chromosomes are compact. Application of the active forces increases *r*_min_, and over a small range of forces *r*_min_ becomes commensurate with 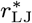 (Figs. 5A–5C). It should be noted that the criterion for solid-like ordering is *independent* of the interaction parameters characterizing the chromosome, and depends only on the magnitude of the force (see SI Appendix, Sec. 10; Fig. S12). If the bonds are rigid, chain segments in the polymer chain, under confinement and active extensile forces, would align exhibiting a nematic order (51). Thus, the mechanisms of our FCC-like ordering and the nematic order in Ref. 51 are qualitatively different. However, both the studies show that active forces induce ordering in chromatin. Additional comparisons between the two models are given in SI Appendix, Sec. 11.

### Studies of activity-induced changes in polymers

The roles of active processes and associated forces in chromosome organization and dynamics have been probed in theoretical studies based on polymer models (6, 50–55). It was shown that athermal random noise, when applied selectively on the active loci, affects the nuclear positioning of the active loci (6, 50, 54). Interestingly, using a combination of analytical theory and simulations (55), it has been shown that a modest difference in activity between active and inactive loci leads to microphase separation (55), a key feature of organization in interphase chromosomes. Using a model that is similar to that used here (54), it has been shown that active forces are important in driving heterochromatin to the periphery of the nucleus, which attests to the dynamic nature of chromatin structures. Typically applying active forces resulted in the increased MSD of the polymer chain (44–46, 55), which would not capture the suppressed chromatin mobility in human cells with active transcription (8, 13, 16). Our minimal model, in the absence of active forces, leads to compact globule that exhibits glass-like dynamics over a long time (10), which may be needed to reproduce the experimental findings. Furthermore, it is likely that explicit inclusion of the precise epigenetic states, as done here for chromosomes 5 and 10, is required for observing the unexpected role of active forces on locus mobilities.

### Effect of active force density

To assess how the force density affects the results, we repeated the simulations for Chr5 by adapting the model such that the active forces are applied to only a fraction of the A-type loci. First, we applied the extensile force dipole on a bond of A-type loci per 20 loci in each gene region (see Fig. 7A for schematic illustration). This implementation corresponds to the RNAPII density of *∼* 0.04 per kb, assuming that there is a linear relation between the force density and the linear RNAPII density. At this density of the active loci, there is no discernible change in the polymer dynamics regardless of the force magnitude (Fig. 7B, inset).

**Fig. 7.**
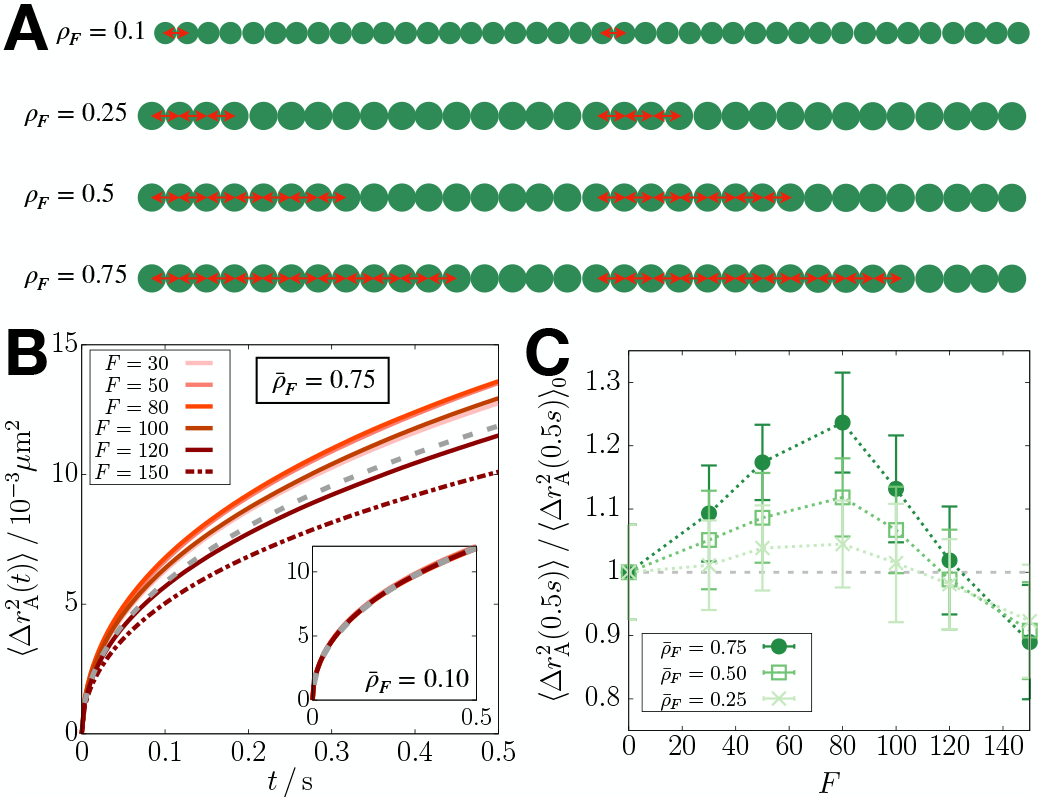
Effect of density of active forces on the mean square displacement. (A) Schematic illustration implementing the active forces at a given density in a single gene region of 40 (*ρ*_*F*_ = 0.1) or 32 (*ρ*_*F*_ *≥* 0.25) loci, where the green circles are the gene loci and the red double-arrows are the active forces. (B) 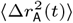 from the simulations at the average force density 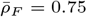 a function of *F*, where the gray dashed line is 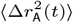 at *F* = 0. The inset shows the results for 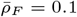. (C) Ratio of the MSD at 0.5 s for the active polymer at given force density to the passive case as a function of *F*. The dotted lines are a guide to the eye.

We then increased the force density gradually to detect any changes in the polymer dynamics. We describe the force density using *ρ*_*F*_ (*i*) = *n*_*F*_ (*i*)*/n*_A_(*i*), where *n*_*F*_ (*i*) is the number of loci subject to active forces in the *i*^th^ gene and *n*_A_(*i*) is the total number of loci in the gene. For instance, the above case in which the forces were applied on a bond every 20 loci corresponds to *ρ*_*F*_ = 0.1, whereas *ρ*_*F*_ = 1 for our original simulations (yielding the results shown in Figs. 2–6). We considered the cases with the average force density, 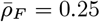, 0.5, and 0.75 (Fig. 7A). At these force densities, the MSD shows a non-monotonic change upon increasing *F*, which differs from that at *ρ*_*F*_ = 1 (Figs. 7B–7C). The MSD increases for *F* ≲ 80 and decreases for *F >* 80. The increase is comparable to that for *ρ*_*F*_ = 1 at *F ≈* 30.

Our model requires a large density of active forces (*ρ*_*F*_ = 1) in order to capture the observed decrease in the chromatin loci mobility upon transcription activation (13, 16). This restriction may have two implications. On the one hand, this might be considered as a limitation of the model. Due to its simplicity, the model might not be able to account for the effect of RNAPII-induced force at a reasonable linear RNAPII density. However, what is more relevant may be the number of RNAPII molecules per unit volume during transcription, which is harder to estimate. Moreover, it is unclear how much active force is transmitted by RNAPII molecules to the gene loci at the resolution used in the model. On the other hand, the simulation results imply that the loci exhibiting mobility decrease upon transcription activation might experience more force than expected from active RNAPII molecules. More precisely, collectively increased bond distances between consecutive gene loci will result in the motility decrease due to the solid-like ordering (see Fig. 5C). Such extra amount of force or extended bonds could arise from ATP-dependent chromatin/nucleosome remodeling that is associated with active transcription (85). Similarly, our model suggests that if there is less amount of force or bond extension in gene regions (*i*.*e*., *ρ*_*F*_ ≲ 0.75), the mobility of the gene loci will increase upon activation without ordering, which can account for the results from Ref. 14 (see the next subsection).

### Connection to other experiments

Gu *et al*. (14) reported that transcription enhances the mobility of specific gene promoters/enhancers, which seems to contradict the results in a related experimental study (13, 16). Gu *et al*. tracked the motion of only a few loci whereas Nagashima *et al*. (13) measured the motility of many loci across the nucleus. In addition, the active genes chosen by Gu *et al*. seem to be outliers in terms of changes in the gene expression level upon cell differentiation (14). Those genes show about 50–100 times difference in their expression levels upon activation or suppression following cell differentiation, although the expression of other genes mostly change by a factor of 4–5. We surmise that the remarkably high level of gene transcription activity results in the mobility changes that are opposite to the global trend for all other genes (8, 13, 16). However, the precise explanation for the differences between the experiments (13, 14, 16) is lacking. Using our results, we interpret, tentatively, that the enhanced motility observed for the activated genes (14) likely corresponds to 0 *≤ F ≤* 30 or 90 *≤ F ≤* 120 at *ρ*_*F*_ = 1 (see Fig. 2C), or 0 *≤ F* ≲ 80 at *ρ*_*F*_ ≲ 0.75 (see Fig. 7C), or possibly to the case where these genes are situated with no other co-activated genes in their proximity (see Discussion below). On the other hand, a majority of the active loci tracked in Nagashima *et al*. (13) likely corresponds to 60 *≤ F≤* 90 at *ρ*_*F*_ = 1.

In another experiment (12), it was shown that acute depletion of cohesin, which mediates the formation of chromatin loops along with CTCFs (33, 39, 86, 87), results in increased motility of nucleosomes. Our polymer simulations capture the enhanced dynamics upon loop loss at *F* = 0. Indeed, when we compare the MSDs between *F* = 80 with loops and *F* = 0 without loops, a near quantitative agreement with the experiment (*∼* 20–25% increase) is obtained. This result implies that the loop loss may lead to the decreased transcription activity of some genes, possibly due to the disruption of interactions between the cognate promoter and enhancer. Although the downregulation of genes was observed upon loop loss (86–88), the overall effect on gene expression is not clearly understood since the loop formation is dynamic (37, 38, 89), and is likely coupled to transcription (90, 91). More precise theory that incorporates the effect of dynamic loops on transcription needs to be constructed using the experiments that measure locus motion and real-time gene expression simultaneously (92).

The physical picture suggested here needs additional validation by future experiments. Direct observation of transient ordering could be challenging in the resolution limit of currently available in imaging methods. Possibly, the live-cell magnetic tweezer technique (64) may be utilized to test our finding. Recent experiments support our conclusions indirectly (30, 93, 94). In particular, Bohrer and Larson (94) highlighted the significant correlation between the spatial proximity and the co-expression of genes by analyzing the DNA and RNA FISH data (95). The role of gene co-activation is implicitly considered in our study because we apply active forces on all the gene-transcribing loci in a given chromosome region. We do not observe the activity-induced decrease in chromatin mobility when only one gene or a few loci are active (see SI Appendix, Sec. 12; Fig. S13). Based on these simulations, we conclude tentatively that motility of loci would increase when only a single gene is transcribed whereas RNAPII-induced forces would suppress chromatin loci mobility when a few genes are co-expressed.

### Limitations

Our model is not without limitations. For instance, explicit effect of transcription factors may bring the theory closer to experiments (96) but will come at the expense of complicated simulations. In addition, our model does not include hydrodynamic effects (51, 97, 98) that contribute significantly to the correlated motion of chromatin observed in the experiments (16). On the experimental side (13), the mobility change might arise due to DNA damage that could be induced by transcriptional inhibition (16, 99). The DNA repair process, which is not considered here, could be an alternate mechanism that could explain the increase in the MSD (16, 100).

For a more realistic picture of gene expression dynamics, our model should be extended to incorporate RNAP-induced force transmission in temporally and/or spatially dependent manner. This extension may also include the implementations of the features at higher resolution, such as the dynamic loop extrusion (35, 37, 38, 91) and the RNAP translocation with nascent RNAs elongated (68). Such a biophysical model would provide clearer pictures on how the gene expression is regulated in conjunction with the dynamically-evolving chromatin topology.

The prediction of transient solid-like ordering might be criticized because currently there is no experimental validation of the proposed mechanism. However, whether chromatin is solid-like (30) or liquid-like (12) continues to be controversial (101). It could adopt both these characteristics depending on length and time scales (102). In light of the unresolved nature of the dynamical state of chromatin, our proposal of the emergence of transient solid-like order is not unreasonable, especially considering that the simulations confirm the non-intuitive experimental observations. Additional experiments are needed to clarify the nature of the chromatin state, which is likely to change depending on the chromatin activity.

## Methods

### Assignment of the monomer type and loop anchors in CCM

We model chromosomes as a copolymer that embodies the epigenetic information. The locus type and loop anchor locations (see Fig. 1) in the CCM are assigned following the procedures given in our previous study (10). The locus type (euchromatin or heterochromatin) is assigned using the epigenetic information from the Broad ChromHMM track (103–105). The first 1–11 states are labeled as active (euchromatin, A) and states 12–15 as inactive (heterochromatin, B). To identify the loop anchors, we use the locations of unique CTCF loops identified by Rao et al. (33). We assign loop anchors to the loci whose genomic regions are closest to the centroids of the experimentally identified anchors. For the results shown above, we considered chromosome 5 (Chr5) in the region from 145.87 to 150.67 Mb from the GM12878 cell line with the resolution of 1.2 kb per locus. The numbers of active and inactive loci are *N*_A_ = 982 and *N*_B_ = 3018, respectively. In this Chr5 region, there are 13 CTCF loops. The simulations were performed using a single copolymer model of Chr5.

### Potential energy function of CCM

The CCM energy function for a chromosome is given by *U*_0_ = *U*_NB_ + *U*_B_ + *U*_L_, where *U*_NB_, *U*_B_, and *U*_L_ represent the energetic contributions of non-bonding interactions, bond stretch, and CTCF-mediated loops, respectively. We use the Lennard-Jones potential for the non-bonding pair interactions, given by 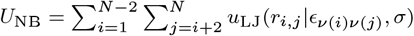, where

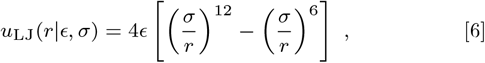

*r*_*i,j*_ = |**r**_*i*_ *−* **r**_*j*_| is the distance between the *i*^*th*^ and *j*^*th*^ loci, and *σ* is the diameter of a locus. The diameter of A and B type loci are the same. Here, *ν*(*i*) is the locus type, A or B, so there are three interaction parameters, *ϵ*_AA_, *ϵ*_BB_, and *ϵ*_AB_. The bond stretch energy is written as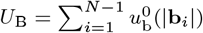, where 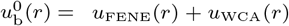 and **b**_*i*_ = **r**_*i*+1_ **r**_*i*_. *u*_FENE_(*r*) is the finite extensible nonlinear elastic (FENE) potential (106, 107),

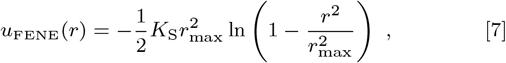

where *r*_max_ is the maximum bond length and *K*_S_ is the FENE spring constant. *u*_WCA_(*r*) is the Weeks-Chandler-Anderson (WCA) potential (108), given by,

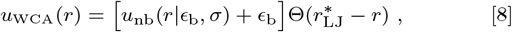

where Θ(*x*) is the Heaviside step function (Θ(*x*) = 1 if *x >* 0 and Θ(*x*) = 0 if *x ≤* 0). The WCA potential is the repulsive tail of the LJ potential, which is shifted and truncated at 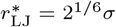, and keeps a pair of adjacent loci from collapsing. Finally, the constrains imposed by the loop anchors is modeled using a harmonic potential,

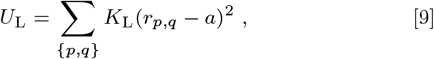

where {*p, q*} is the set of indices of the loop anchors, *a* is the equilibrium bond length between the pair of loop anchors, and *K*_L_ is the harmonic spring constant.

The values of the parameters in *U*_B_ and *U*_L_ are *K*_S_ = 30*k*_*B*_*T/σ*^2^, *r*_max_ = 1.5*σ, ϵ*_b_ = 1.0*k*_*B*_*T, K*_L_ = 300*k*_*B*_*T/σ*^2^, and *a* = 1.13*σ*. With these parameter values, the equilibrium bond length is given by *b*_0_ = 0.96*σ* at which 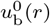 is minimized. For the energetic parameters in *U*_NB_, we used *ϵ*_AA_ = *ϵ*_BB_ = 2.4*k*_*B*_*T* and 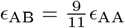. The previous study showed that the CCM with this parameter set reproduces the Hi-C inferred contact maps for chromosomes 5 and 10 well (10). In particular, compartment features on *∼* 5 Mb scale and topologically associating domains (TADs) on *∼* 0.5 Mb are reproduced using the CCM simulations. In addition, the CCM simulations reproduced the dynamic properties that agree with experimental results.

### Simulation details with active force

We performed Brownian dynamics simulations by integrating the following equation of motion, which for the *i*^th^ locus is given by,

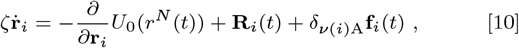

where *ζ* is the friction coefficient and *U*_0_(*r*^*N*^ (*t*)) is the potential energy of the CCM polymer chain with the configuration, *r*^*N*^ (*t*) = {**r**_1_(*t*),, **r**_*N*_ (*t*)}). The Gaussian white noise, **R**_*i*_(*t*) satisfies ⟨ **R**_*i*_(*t*) **R**_*j*_ (*t*^*′*^) ⟩ = 6*ζk*_*B*_*Tδ*_*ij*_ *δ*(*t − t*^*′*^), where *k*_*B*_ is the Boltz-mann constant and *T* is temperature. The Kronecker delta, *δ*_*ν*(*i*)A_, in the last term of Eq. 10 ensures that **f**_*i*_(*t*) acts only on the A-type loci. The exact form of **f**_*i*_(*t*) is,

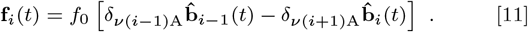

The time step for integrating Eq. 10 was chosen as *δt* = 10^*−*4^*τ*_*B*_, where *τ*_*B*_ is the Brownian time, defined by *τ*_*B*_ = *σ*^2^*/D*_0_. Here, *D*_0_ = *k*_*B*_*T/ζ* is the diffusion coefficient of a single locus. Using the Stokes-Einstein relation, *D*_0_ = *k*_*B*_*T/*6*πηR*, where *η* is the viscosity of the medium and *R* is the radius of a locus, we can evaluate the diffusion coefficient and the simulation time step in real units. We choose *η* = 0.89 *×* 10^*−*3^ Pa · s from the viscosity of water at 25°C *≈* and *R σ/*2. We take *σ* = 70 nm from an approximate mean of the lower and upper bounds for the size of 1.2 kb of chromatin including six nucleosomes and linker DNAs, which are estimated as 20 nm and 130 nm, respectively (10). Hence, in real time and length units, 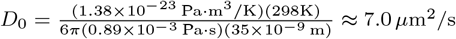 and *τ*_*B*_ *≈* 0.7 ms.

For the most part, the results of the simulations are reported in the reduced units. The energy parameters are given in units of *k*_*B*_*T*. Thus, the fundamental energy unit is *ϵ* = *k*_*B*_*T*, which means that the reduced temperature becomes *T*^*\**^ = *Tk*_*B*_*/ϵ* = 1. The fundamental length and time units are *σ* and *τ* = (*mσ*^2^*/ϵ*)^1*/*2^, respectively, where *m* is the mass of a single locus. With an estimate of 260 kDa per nucleosome with 200 bps of dsDNA, we obtain *τ ≈* 55.5 ns in real units so the time step is given by *δt ≈* 1.26*τ*. The magnitude of the active force is *F* = *f*_0_*σ/ϵ* = *f*_0_*σ/k*_*B*_*T*. For instance, *F* = 80 corresponds to 3 to 16 pN in real units, based on the lower and upper bounds of *σ* specified above (20 nm to 130 nm).

Each simulation starts from a collapsed globule configuration that is obtained from the equilibration of an extended polymer chain using the low-friction Langevin thermostat at *T*^*\**^ = 1.0 in the absence of active force. We propagate the collapsed configuration with active force for 10^6^*δt* until the active polymer reaches a steady state where the radius of gyration and potential energy does not increase further. Subsequently, we run the simulation for additional 2 *×* 10^8^*δt* and generate five independent trajectories to obtain the statistics needed to calculate the quantities of interest. All the simulations were performed using LAMMPS (109).

### Inherent structure for the active polymer

Following the concept introduced by Stillinger and Weber (73), we investigated the *inherent structure* for each locus type in the CCM. The inherent structure is the ideal structure in molecular liquid which is preferred by the geometrical packing of particles, so it is the arrangement of particles expected by removing thermal excitations. Practically, the inherent structure is determined by minimizing the energy of the system using the steepest descent method (110). This procedure, however, is not directly applicable to our system because it involves the active force that is not derived from a given potential energy. Instead, we perform the quench of the CCM with the active force to low temperature. In a previous study, it was shown that quenching the monodisperse colloidal liquid generates a stable BCC (body-centered cubic) crystal, which cannot be easily obtained by the standard steepest descent (74).

To quench the system to a low temperature, we took the configuration from *T* = 298*K* (*T*^*\**^ = 1.0) and ran Brownian dynamics simulation at *T ≈* 3 K (*T*^*\**^ = 0.01). The Brownian time and simulation time step are scaled accordingly such that 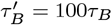 and 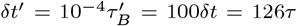. The quench simulation was run *∼* for 10^6^*δt*^*′*^ during which the system energy was minimized until it reached a plateau value within ∼10^5^*δt*^*′*^. We computed the radial distribution functions, *g*_AA_(*r*) and *g*_BB_(*r*), from the quenched configurations.

### Bond-orientational order parameter

Following Ref. 75, we computed the bond-orientational order (BOO) parameter for each individual locus. The BOO with *l*-fold symmetry for the *i*^th^ locus is defined as,

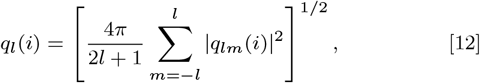

where *q*_*lm*_(*i*) is the average of the spherical harmonics, *Y*_*lm*_, for the bond angles formed between the *i*^th^ locus and its nearest neighbors,

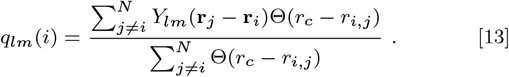

We use *r*_*c*_ = 1.4*σ*, where the radial distribution function has the local minimum after the peak corresponding to the first coordination shell, as the cutoff pair distance for the nearest neighbors. The probability distribution and average of BOO parameter were computed depending on the locus type such that

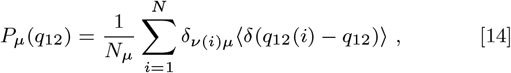

and

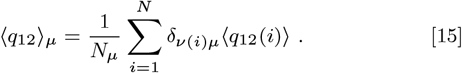

The distributions, *P*_A_(*q*_12_) and *P*_B_(*q*_12_), for different values of *F* are shown in Figs. S4E and S4F.

### Effective potential energy

Since the active force is not stochastic, it may be treated as a pseudo-conservative force, which contributes to potential energy. In particular, we define the effective potential energy,

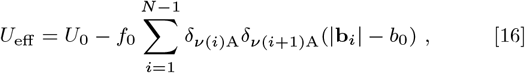

where the second term, denoted by *U*_*a*_, represents the work due to the active force on the A-A bonds. It can be verified that (*∂/∂***r**_*i*_)*U*_a_ = − *δ*_*ν* (*i*)A_**f**_*i*_, so the equation of motion (Eq. 10) can be rewritten as,

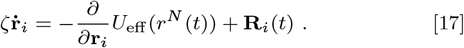

The minus sign in *U*_*a*_ indicates that this potential prefers bond extension. Note that *U*_*a*_ only affects the bond potential energy of the _b_ bonds. It is consistent with the effective bond potential, 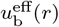 in Eq. 5, with 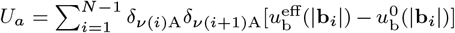. In Fig. 5D, we showed the mean and variance of the effective potential energy for the A loci (*E*_eff,A_) as a function of *F*. *E*_eff,A_ is given by the sum of all the pairwise interactions involving the A loci, or the total effective potential energy excluding the contributions of B-B pairs,

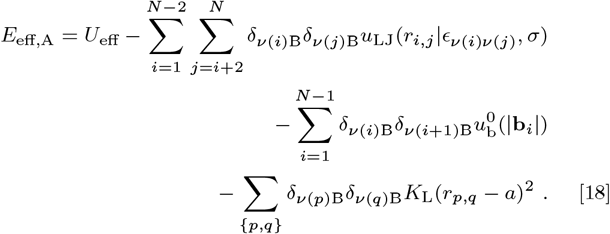

## ACKNOWLEDGMENTS

We thank Alexandra Zidovska, Bin Zhang, Debayan Chakraborty, Xin Li, and Davin Jeong for useful discussions. We thank the National Science Foundation (CHE19-00093) and the Welch Foundation through the Collie-Welch Chair (F-0019) for support.

