## Supplementary Information for "Transcription-induced active forces suppress chromatin motion"

#### This PDF file includes:

Supplementary text
Figs. S1 to S13
Legends for Movies S1 to S2
SI References

#### Other supplementary materials for this manuscript include the following:

Movies S1 to S2

### Supporting Information Text

#### 1. Biological relevance of transcription-induced active force in the model

We assume that the active force is applied along the chromatin chain in order to model transcriptional elongation. Although modeling the complex process of transcription using mechanical force is an oversimplification, we argue that it is reasonable. Single-molecule experiments have shown that RNA polymerase II (RNAPII) moves along DNA exerting large forces (up to 45 pN or more) during transcription (S1). The helicase activity of RNAP, resulting in base pair opening, implies that the forces are exerted along the DNA bonds. Our simple active chromatin model provides insights into the dynamics of chromosomes in response to RNAP binding, which are difficult to obtain using experiments alone.

The locus resolution, 1.2 kb $\sim$ 6 nucleosomes connected by linker DNAs, is smaller than the size of a typical gene, which is about 27 kb in human chromosomes (S2). At this resolution, the intergenic regions that correspond to inactive (B-type) loci are well separated from the gene-transcribing regions (A-type loci) in our model. Each gene body, represented by multiple A-type loci, is presumed to contain a few actively transcribing RNAPII, as observed in the active burst phase of human genes (S3).

Is it reasonable to assume that the forces arising from the activity of RNAP in the gene-transcribing region propagate through other loci in the gene? We believe it is because the time scale for force propagation across the gene loci is expected to be less than the transcription time. Indeed, the time scale for force propagation in our monomer length scale is roughly 1 ns *at most*, based on the Young's modulus and mass density of DNA, which is smaller than our simulation time unit ($\sim$ 70 ns; see Methods in the main text). So the locally applied force by RNAP is felt outside the gene-transcribing regions (*i.e.*, throughout a given A-type locus).

As described in the main text (see Fig. 1D), the active force is exerted on each bond of the A-type loci in the form of a divergent or dipolar pair parallel to the bond vector, the magnitude of which is in accord with optical tweezer experiments (S1). Such a force dipole should increase the distance between a bonded active locus pair. The increase in the bond distance is accompanied by an increase in the effective excluded volume of the A loci, potentially due to the local nucleosome motion enhanced by the tension forces propagated from actively transcribing RNAPs (see Fig. 1D). In the overdamped regime (as implemented in Eq. 10), the propagation of $f_{\mathrm{RNAP}} \approx 10$ pN would make the nucleosomes diffuse with the effective speed of $v = \frac{f_{\mathrm{RNAP}}}{\zeta_{\mathrm{nuc}}} \approx 5\,\mathrm{nm}$ per simulation time step, where $\zeta_{\mathrm{nuc}}$ is the friction coefficient for a nucleosome-sized particle in water. The locally activated motion of nucleosomes would increase the bond length between the monomers by inducing the volume exclusion. The magnitude of the volume-excluding repulsion should be comparable to that of the tensional force on DNA exerted by RNAP. The typical estimate of $\approx 16$ pN at $F = 80$ used in our simulations is reasonable.

#### 2. Relevant time range of the experimental MSD data for comparison

Nagashima *et al.* tracked the motions of individual nucleosomes in live human RPE-1 cells and reported the mean square displacements (MSDs) of the nucleosomes with or without active transcription (S4). They found that the MSD as a function of time reached a plateau within about 3 seconds, as shown in Fig. 1A. The saturation of the MSD curve is attributed to the constraint imposed on the nucleosome motion within the densely packed environment of chromatin. In the MSD curve, we find that there are two different scaling regimes depending on the time scale. For the control case (black color), the scaling exponent is about 0.5 for $t \leq 0.5\,\mathrm{s}$ whereas 0.23 for $t > 0.5\,\mathrm{s}$. The cases treated with the transcription inhibitors, $\alpha$-amanitin ($\alpha$-AM) and 5,6-dichloro-1-$\beta$-D-ribofuranosylbenzimidazole (DRB), show qualitatively the same scaling behavior. Since the scaling behavior for $t \leq 0.5\,\mathrm{s}$ is more relevant to the polymer diffusion, we considered the data only for $t \leq 0.5\,\mathrm{s}$ to compare with our simulation results.

### 3. Contributions of A- and B-type loci to dynamic properties

In the main text (Fig. 2B), we compared the difference between the passive and active cases of our simulations for a particular type of loci with the experimental results. Nagashima *et al.* analyzed the MSDs based on the locations within the cell nucleus, specifying the data measured from either the interior or periphery of the nucleus (S4). We assumed that the experimental results for the interior (periphery) can be approximated as for euchromatin (heterochromatin). For both A and B loci, the change in MSD upon treating the inhibitor is qualitatively comparable to the simulation results (Figs. 2A and S1A). For a quantitative comparison, we calculated the relative increase in $\langle \Delta r_{\mathrm{A}}^2(t)\rangle$ and $\langle \Delta r_{\mathrm{B}}^2(t)\rangle$ upon turning off the activity, using

$$\Delta\overline{\mathrm{MSD}}_{\mathrm{exp}}^{\mu} = \frac{1}{t_{\mathrm{diff}}}\int_0^{t_{\mathrm{diff}}} dt \frac{\langle \Delta r_\mu^2(t)\rangle_{\mathrm{inhibited}} - \langle \Delta r_\mu^2(t)\rangle_{\mathrm{control}}}{\langle \Delta r_\mu^2(t)\rangle_{\mathrm{control}}} \ , \tag{S1}$$

for the experiment and

$$\Delta\overline{\mathrm{MSD}}_{\mathrm{sim}}^{\mu} = \frac{1}{t_{\mathrm{diff}}}\int_0^{t_{\mathrm{diff}}} dt \frac{\langle \Delta r_\mu^2(t)\rangle_{F=0} - \langle \Delta r_\mu^2(t)\rangle_{F=80}}{\langle \Delta r_\mu^2(t)\rangle_{F=80}} \ , \tag{S2}$$

for the simulation, where $t_{\mathrm{diff}} = 0.5\,\mathrm{s}$.

We found that the comparison between $\Delta\overline{\mathrm{MSD}}_{\mathrm{exp}}$ and $\Delta\overline{\mathrm{MSD}}_{\mathrm{sim}}$ is more quantitative for the B loci than the A loci (Fig. 2B, main text). The quantitative difference could arise because every interior measurement does not necessarily correspond to the euchromatin whereas the periphery measurements are mostly for the heterochromatin. It is notable that the comparison between simulations and experiments is qualitatively similar for $F$ in the range, $60 \leq F \leq 90$ (Fig. S2). If we compare the overall MSD for all the loci, $\langle \Delta r^2(t)\rangle = [N_{\mathrm{A}}\langle \Delta r_{\mathrm{A}}^2(t)\rangle + N_{\mathrm{B}}\langle \Delta r_{\mathrm{A}}^2(t)\rangle]/N$ with the experimental results (Fig. S1B), we find that $\Delta\overline{\mathrm{MSD}}_{\mathrm{sim}}$ quantitatively agrees with $\Delta\overline{\mathrm{MSD}}_{\mathrm{exp}}$ (Fig. S1C). Note that the overall MSD, $\langle \Delta r^2(t)\rangle$, at a given lag time (*i.e.*, $t = 100\tau_B \approx 0.07\,\mathrm{s}$) reflects the non-monotonic change of $\langle \Delta r_{\mathrm{A}}^2(t)\rangle$ upon increasing $F$ from 0 to 120 (Fig. S1D).

We also decomposed $\langle F_s(k_{\max}, t)\rangle$ and $\chi_4$, shown in Fig. 3 of the main text, into contributions from the A and B loci separately. For the $\mu$-type loci, we define

$$F_s^\mu(|\mathbf{k}|, t) = \frac{1}{N_\mu}\sum_{i=1}^{N} \delta_{\nu(i)\mu} e^{i\mathbf{k}\cdot[\mathbf{r}_j(t) - \mathbf{r}_j(0)]} \ , \tag{S3}$$

and

$$\chi_4^\mu(t) = N_\mu\left[\left\langle F_s^\mu(k_{\max}, t)^2\right\rangle - \langle F_s^\mu(k_{\max}, t)\rangle^2\right] \ , \tag{S4}$$

where $k_{\max} = 2\pi/\sigma$. Figure S3A compares $\langle F_s^{\mathrm{A}}(k_{\max}, t)\rangle$ and $\langle F_s^{\mathrm{B}}(k_{\max}, t)\rangle$. For $F = 0$, $\langle F_s^{\mathrm{A}}(k_{\max}, t)\rangle$ decays faster than $\langle F_s^{\mathrm{B}}(k_{\max}, t)\rangle$, whereas $\langle F_s^{\mathrm{A}}(k_{\max}, t)\rangle$ decays slower at $F = 80$. Remarkably, $\langle F_s^{\mathrm{A}}(k_{\max}, t)\rangle$ at $F = 80$ exhibits a shoulder prior to the structural relaxation. This shoulder, starting near $\langle F_s^{\mathrm{A}}(k_{\max}, t)\rangle = 0.7$ and $t < \tau_B$, arises due to the enhanced caging by neighboring particles around the probe, which results from the collective solid-like ordering. The Fourier transform of the scattering function (Fig. S3A, inset) illustrates the increase in the slowly-decaying low-frequency modes, which correspond to the collective motions of the ordered particles, whereas the fast-decaying high-frequency modes are rather random motions. The similar trend is also reflected in $\chi_4^{\mathrm{A}}(t)$ and $\chi_4^{\mathrm{B}}(t)$, as shown in Fig. S3C. At $F = 0$, the peak of $\chi_4^{\mathrm{A}}(t)$ appears earlier than $\chi_4^{\mathrm{B}}(t)$, albeit with a smaller amplitude. In contrast, at $F = 80$, $\chi_4^{\mathrm{A}}(t)$ exhibits a larger peak which appears at a later time than $\chi_4^{\mathrm{B}}(t)$, suggestive of large-scale coherent motions. In Fig. S3B, the plots of $\tau_\alpha^{\mathrm{A}}$ and $\tau_\alpha^{\mathrm{B}}$, the relaxation times determined from $\langle F_s^{\mathrm{A}}(k_{\max}, t)\rangle$ and $\langle F_s^{\mathrm{B}}(k_{\max}, t)\rangle$, respectively, indicate that the retardation of the active polymer chain dynamics compared to the passive case is predominantly driven by the changes in the dynamics of the A loci. In accord with this observation, the increase in dynamic heterogeneity at $F > 0$ is also originated from the A loci rather than the B loci (Fig. S3D).

**Alignment of the sampled configurations.** When analyzing the simulation data to compute the dynamic properties, we align the whole polymer with respect to the initial configuration to remove the global translational and rotational drifts. Specifically, at each time frame in a given trajectory, we shift the polymer configuration to minimize the root mean square displacement from the initial configuration. As a result, $\langle \Delta r_\mu^2(t) \rangle$ measures the internal diffusion of individual loci in the reference frame of the center of mass of the system.

### 4. Bending angle distribution

The extensile active force applied to each A-A bond could induce a net perpendicular force to the chain contour for the A loci, as implied in Fig. 1E in the main text. The net force on the $i^{\text{th}}$ locus, given in Eq. 11 if the $(i-1)^{\text{th}}$ and $(i+1)^{\text{th}}$ are also A-type, becomes $\mathbf{f}_i(t) = -f_0 \boldsymbol{\kappa}_i(t)$ where $\boldsymbol{\kappa}_i(t) = \hat{\mathbf{b}}_i(t) - \hat{\mathbf{b}}_{i-1}(t)$ is the curvature vector at the $i^{\text{th}}$ point of the discrete chain connected by the normal bond vectors. Hence, the applied force is should decrease the local bending angles for the A-type loci. To probe the effect of the force on the curvature, we calculated the distribution of the angle formed between two consecutive A-A bonds, defined as,

$$P(\theta_{\mathrm{A}}) = \frac{\sum_{i=2}^{N-1} \delta_{\nu(i-1)\mathrm{A}} \delta_{\nu(i)\mathrm{A}} \delta_{\nu(i+1)\mathrm{A}} \langle \delta(\cos^{-1}(\hat{\mathbf{b}}_i \cdot \hat{\mathbf{b}}_{i+1}) - 180^\circ + \theta_{\mathrm{A}}) \rangle}{\sin\theta_{\mathrm{A}} \sum_{i=2}^{N-1} \delta_{\nu(i-1)\mathrm{A}} \delta_{\nu(i)\mathrm{A}} \delta_{\nu(i+1)\mathrm{A}}} . \qquad \text{[S5]}$$

Upon increasing the activity, the distribution becomes broader as the peak at $\theta_{\mathrm{A}} = 71^\circ$ shifts to the left (Fig. S5, top), indicating that the bending angle tends to decrease due to the active force. Remarkably, $P(\theta_{\mathrm{A}})$ at $F = 80$ shows more pronounced peaks at $\theta_{\mathrm{A}} = 60^\circ$, $120^\circ$, and $180^\circ$, along with another one at $\theta_{\mathrm{A}} = 90^\circ$, which reflect the structural transition involving the transient FCC-like order. The less pronounced peak near $\theta_{\mathrm{A}} = 147^\circ$ is due to the hexagonal closed packing (HCP) geometry that arises at the surface of the polymer globule (S5). In contrast, the bending angle distribution for the B loci, $P(\theta_{\mathrm{B}})$, similarly defined as in Eq. S5, does not change upon increasing $F$ (Fig. S5, bottom).

### 5. Correlation time of bond-orientational order parameter

We calculated the temporal correlation in $q_{12}$ for a given locus type by using the time correlation function,

$$C_{q_{12}}^{\mu}(t) = \frac{\langle \delta q_{12}(t) \delta q_{12}(0) \rangle_\mu}{\langle (\delta q_{12})^2 \rangle_\mu} = \frac{1}{\langle (\delta q_{12})^2 \rangle_\mu} \left[ \frac{1}{N_\mu} \sum_{i=1}^{N} \delta_{\nu(i)\mu} \langle q_{12}(i;t) q_{12}(i;0) \rangle - \langle q_{12} \rangle_\mu^2 \right] . \qquad \text{[S6]}$$

In Fig. S6B, $C_{q_{12}}^{\mathrm{A}}(t)$ and $C_{q_{12}}^{\mathrm{B}}(t)$ are shown for five independent simulation trajectories at $F = 80$. Due to activity-induced ordering in the A loci, $C_{q_{12}}^{\mathrm{A}}(t)$ decays over longer time scale than $C_{q_{12}}^{\mathrm{B}}(t)$. Nevertheless, a majority of the order decorrelates in less than $0.2\tau_B$ (Fig. S6B, inset). The maximum time over which $C_{q_{12}}^{\mathrm{A}}(t)$ decays is $1,000\tau_B \approx 0.7\,s$. The transient nature of the order is also visually discernible from a trajectory at $F = 80$, if we mark the loci whose $q_{12}$ is larger than a threshold at a given time. The threshold is chosen to be $q_{12} = 0.45$, which is approximately the midpoint between the liquid ($q_{12} \approx 0.3$) and the FCC solid ($q_{12} \approx 0.6$). The simulation snapshot with this labeling illustrates that only subpopulations of the A loci exhibit the order (Fig. S6C, left). This ordering fluctuates from trajectory to trajectory. In contrast, the quenched active polymer shows a larger cluster of FCC-like ordering that stays indefinitely (Fig. S6C, right). The movies for the trajectories including the snapshots shown in Fig. S6C are given in Movies S1 and S2.

### 6. FCC-like ordering does not alter the chromosome conformation

Does the FCC-like order induced by the active force affect the structure of the folded chromosome? To answer this question, we computed $P(s)$, the average contact probability at a given genomic pair distance,

$s$, using

$$P(s) = \frac{1}{N-s}\sum_{i=1}^{N-1}\sum_{j=i+1}^{N}\delta_{|i-j|,s}C_{i,j}\ , \tag{S7}$$

where $C_{i,j}$ is the contact matrix from either the Hi-C experiment or the simulation. In our previous study (S6), we demonstrated that the CCM quantitatively reproduces the $s$-dependence of the pair contact probability, especially the power-law scaling exponent in $P(s)$. In Fig. S7A, $P(s)$ for the CCM with $F = 80$ is nearly identical to that for $F = 0$, showing reasonable agreement with the Hi-C result by capturing the two distinct scaling exponents in the power-law decay ($\sim s^{-0.8}$ for $s \lesssim 0.5$ Mbps and $\sim s^{-1.5}$ for $s > 0.5$ Mbps). We also calculated the average contact probability for a given pair type, using

$$P_{\mu\gamma}(s) = \frac{\sum_{i=1}^{N-1}\sum_{j=i+1}^{N}\delta_{\nu(i)\mu}\delta_{\nu(j)\gamma}\delta_{|i-j|,s}C_{i,j}}{\sum_{i=1}^{N-1}\sum_{j=i+1}^{N}\delta_{\nu(i)\mu}\delta_{\nu(j)\gamma}\delta_{|i-j|,s}}\ , \tag{S8}$$

where $\mu$ and $\gamma$ is either A or B. As shown in Fig. S7B, there is no significant change in the $s$-dependence of $P_{\mu\gamma}(s)$ upon increasing $F$ from 0 to 80.

For a further comparison between the 3D structures, we plotted the mean distance matrices, $\bar{r}_{i,j}$, at $F = 0$ and $F = 80$ in Fig. S7C. Here, the bar in $\bar{r}_{i,j}$ denotes the average over a single trajectory, and thus we compare the averaged 3D structures that are propagated from the same initial configuration but with different $F$. The visual inspection suggests that there is no significant difference between the distance matrices. The evaluation of the relative mean absolute error (RMAE), defined as

$$\mathrm{RMAE} = \frac{2}{N(N+1)}\sum_{i=1}^{N}\sum_{j=i}^{N}\frac{\bar{r}_{i,j}(F=0) - \bar{r}_{i,j}(F=80)}{\bar{r}_{i,j}(F=0)}\ , \tag{S9}$$

yields $\sim 10\%$, which also indicates quantitative similarity. In Fig. S7D, the calculated and Hi-C contact maps are compared. The simulated contact maps, both at $F = 0$ and $F = 80$, faithfully capture the large-scale patterns in the Hi-C contact map (Fig. S7D), although there are quantitative deviations in the A/B segregation on small length scale (Fig. S7E). Such deviations could arise mainly because of the minimal features of our polymer model as well as the finite-size effect (see the caption of Fig. S7E for more detail). Nevertheless, the predicted contact map at $F = 80$ is essentially the same as the one at $F = 0$. Therefore, the activity-induced ordering does not alter the overall chromosome organization significantly.

### 7. Dynamic properties of a chromosome 10 segment

We also simulated a 4.8-Mbp segment of chromosome 10 (Chr10), which has more A-type loci than the 4.8-Mbp segment of Chr5 (Fig. S9A); $N_{\mathrm{A}}/N = 0.58$ for Chr10 whereas 0.25 for Chr5. We calculated the dynamical properties from the simulations with active force to compare with the findings for Chr5 presented in the main text. In Fig. S9B, the MSDs for different $F$ values are shown in log scale. The dynamics becomes slower upon increasing $F$ and the mobility suppression is maximized at $F = 80$, which follows the same trend found for Chr5 (Fig. S9C). Notably, the extent of the suppression is larger while the scaling exponent of the MSD for $F = 80$ is smaller than for Chr5 ($\alpha = 0.40$ versus 0.43). Similarly, the plots of $\langle F_s(k_{\max}, t)\rangle$ and $\chi_4(t)$ show the trends that are consistent with those for Chr5 (Figs. S9D and S9E); the relaxation is slower and the dynamic heterogeneity is larger than in Chr5. The change due to active force in Chr10 is larger than Chr5, which is a consequence of the higher fraction of A-type loci in Chr10 compared to Chr5. The radial distribution function for A-A pairs, $g_{\mathrm{AA}}(r)$, and the distribution of the BOO parameter for the A loci, $P_{\mathrm{A}}(q_{12})$, show the same trends as those for Chr5 shown in Figs. 4A and 4B of the main text. Overall, the qualitative results are similar between Chr5 and Chr10, which shows that ordering of the active loci for a range of $F$ is a robust finding.

### 8. Random copolymer with $F \neq 0$ does not exhibit fluid-to-FCC transition

We also investigated the effect of $F$ on a copolymer chain with random sequence by shuffling the loci in Chr5 without changing the number of A or B type loci. We performed Brownian dynamics simulations with the active force applied in the same manner as in the wild type (WT) Chr5. This randomized chain does not undergo microphase separation like the WT sequence, whose A/B sequence reflects the genetic activity of an actual chromosome (Chr5 and Chr10). As the favorable interactions between the monomers of the same type is less likely in a random sequence, the dynamics of the randomized chain is moderately faster than the WT, as shown in Fig. S10A. Nonetheless, the dynamics is glass-like (Figs. S10B-S10C), as shown previously (S6). Upon increase of $F$, as shown in Fig. 6A, the MSD for the random sequence decreases by less than 2% only if $F$ exceeds 80. The decomposition of the MSD into A and B contributions shows that the slightly reduced dynamics is driven by the B-type monomers (Fig. 6A). In contrast, the movement of the A-type is enhanced due to the active force. The distributions of the BOO parameters, $P_{\mathrm{A}}(q_{12})$ and $P_{\mathrm{B}}(q_{12})$, confirm that the reduced dynamics for $F > 80$ is not accompanied by the FCC-lattice formation, as found in the WT sequence (Figs. 6B and S10D).

### 9. Effect of removal of loop anchors on the dynamics

We tested if the CTCF-mediated loops contribute to the change in the active loci dynamics observed when the extent of activity is varied. We ran Brownian dynamics simulations of the single CCM chain (Chr5) with all the loops removed (*i.e.*, there is no harmonic potential applied between the locus pairs identified as the CTCF loop anchors). It should be noted that CTCF loop loss eliminates TAD-like structures but enhances compartment formation (microphase separation between A and B type loci). Other simulation details are the same as described above.

For the CCM without the loops, the overall MSD also decreases to almost the same extent as with the loops, upon increasing $F$ from 0 to 80 (Fig. S11A). There is a modest increase in the MSD at $F = 0$ compared to the case with the loops, which is consistent with the experimental observation for the dynamic change upon the CTCF-cohesin loop loss (S7). The MSD for the A-type loci decreases modestly, with the scaling exponent decreasing from from 0.38 to 0.36 (Fig. S11B). The change in the MSD for the B-type loci upon removing the loops is relatively minor (Fig. S11C). We also found that the change in $\langle F_s^{\mu}(k_{\max}, t)\rangle$ and $\chi_4^{\mu}(t)$ due to the loop disruption is not significant either at $F = 0$ or $F = 80$ (Figs. S11D-S11E). Interestingly, although the loop removal leads to more suppressed A-loci dynamics at $F = 80$, the distribution of $q_{12}$ for the A-type loci is nearly identical to that with the loops (Fig. S11F). The conservation of $P_{\mathrm{A}}(q_{12})$ suggests that the transition between disordered and ordered states occurs more frequently, which makes long-time scale motions slower, while the coexistence point remains the same at a given level activity. Therefore, the contribution of the loop constraints to the A-loci dynamics is not so pronounced as that of the morphological transition induced by the active force.

### 10. Effects of variations in non-bonding interactions

As specified in a previous section, we applied the condition of $\epsilon_{\mathrm{AA}} = \epsilon_{\mathrm{BB}}$ to the non-bonding interaction parameters for the CCM simulations. This simple assumption was sufficient to demonstrate the CCM's capability of capturing experimental results faithfully (S6). However, this assumption may not always hold because heterochromatin (B-type) is considered to have more compact nature than euchromatin (A-type). To test if the differentiated A-A and B-B interactions affect our results, we performed additional simulations for the Chr5 segment with $\epsilon_{\mathrm{AA}} < \epsilon_{\mathrm{BB}}$, which represents a larger extent of compaction for the B-type loci than the A-type. Specifically, we set $\epsilon_{\mathrm{AA}} = 2.4k_BT$ and $\epsilon_{\mathrm{BB}} = 2.6k_BT$, whereas we used the same A-B interaction parameter, $\epsilon_{\mathrm{AB}} = \frac{9}{11}\epsilon_{\mathrm{AA}} = 1.96k_BT$. These interaction parameters still ensure microphase separation between A and B loci, as determined by the Flory-Huggins theory (S8, S9). Other simulation details are the same as described above.

In Fig. S12A, the MSD for the A-type loci decreases with $F = 80$ and still captures the experimental results qualitatively (*cf.* Fig. 2A, main text). In Fig. S12B, the MSD for the A-type at $t = 10^3\tau_B \approx 0.7s$

shows the dynamic re-entrance with respect to $F$ whereas the change in the MSD for the B-type is relatively insignificant, which is similar to the result in Fig. 2C. In addition, the structural quantities, $g_{\mathrm{AA}}(r)$ (Fig. S12C) and $P_{\mathrm{A}}(q_{12})$ (Fig. S12D), do not change from the original results shown in Figs. 4A and 4B. The FCC-like arrangement of the active loci is visually noticeable from the simulation snapshot in Fig. S12C. Therefore, the $F$-induced dynamical slowdown and transient ordering of the A loci are not affected by increasing $\epsilon_{\mathrm{BB}}$ relative to $\epsilon_{\mathrm{AA}}$. We surmise that the most important conclusion that the locus dynamics is suppressed during transcription, which is in accord with experiments, is robust as shown in Fig. S12.

### 11. Comparisons with a previous study (S10)

Saintillan, Shelley, and Zidovska (SSZ) studied the effect of extensile active force on the chromosome dynamics using a polymer model (S10). Although both the present work and the SSZ study show that active forces induce order in chromatin, there are a few differences as well. (1) In the SSZ model the bond length is fixed. Here, it is the activity-induced (in a narrow range of $F$) increase in the bond length that produces transient solid-like ordering by making the bond length commensurate with the non-bonding attractive interaction (see the main text). In the SSZ study, nematic order, which emerges in the presence of activity, increases monotonically as $F$ increases, whereas in our model the emergence of solid-like order is non-monotonic. (2) The SSZ simulations were performed by confining the polymer chain, which likely produces alignment. Our simulations do not confine the single chain, and hence the mechanism of FCC-like ordering is different from the formation of nematic phase. (3) Activity in the SSZ arises due to fluid flow that is coupled to motion of the beads on the chain, which is an insightful model for probing the effect of active forces on a polymer. In the present work, we envision that RNAPII exerts forces during transcription. (4) In the unconfined case, the SSZ model predicts coil-stretch transition in the absence of activity (see Fig. S1 in SSZ). In our model, at low $F$ the dynamics exhibits glass-like behavior with no ordering. Despite these differences the formation of ordered structures promoted by active forces is a common theme in both the studies.

### 12. Effect of gene co-activation on the dynamics

A recent study (S11) showed that there is a significant correlation between spatial proximity of genes and their co-expression. Our study provides some insight into the underlying mechanism for the spatial coupling of co-activated genes since active forces are applied to all the active genes in the simulated Chr5 region. It is worth examining the dynamical consequences when only one gene is active and others are turned off. To address this issue, we repeated the simulations by applying the active forces only to the loci corresponding to a particular single gene. We considered the *TCOF1* gene, which is located at 149,737,202–149,779,871 bp on Chr5 (see Fig. S13A). The *TCOF1* gene is about 43 kb long and corresponds to 37 loci ($3223 \leq i \leq 3259$) in the CCM polymer chain ($N = 4000$). Because this gene is not insulated by the CTCF loops, it is potentially less affected by the loops. Application of the active force on a single gene results in enhanced mobility of the gene loci (Fig. S13B), whereas the overall MSD for all the A loci does not show a noticeable change (Fig. S13B, inset). We repeated the simulations by exerting the active forces only to the first two consecutive A-A bonds in the gene, *i.e.*, three A-type loci ($i = 3223, 3224, 3225$). The active forces on the three loci give some changes in the MSD of those loci, but the variations are so large that the overall change upon increasing the activity is not significant (Figs. S13C and S13D). Fig. S13D shows that the MSD for the gene loci increases monotonically as $F$ is increased when only one gene is transcribed. Therefore, it is reasonable to suggest that activity-induced mobility suppression requires co-activation of genes in the given chromosome region.

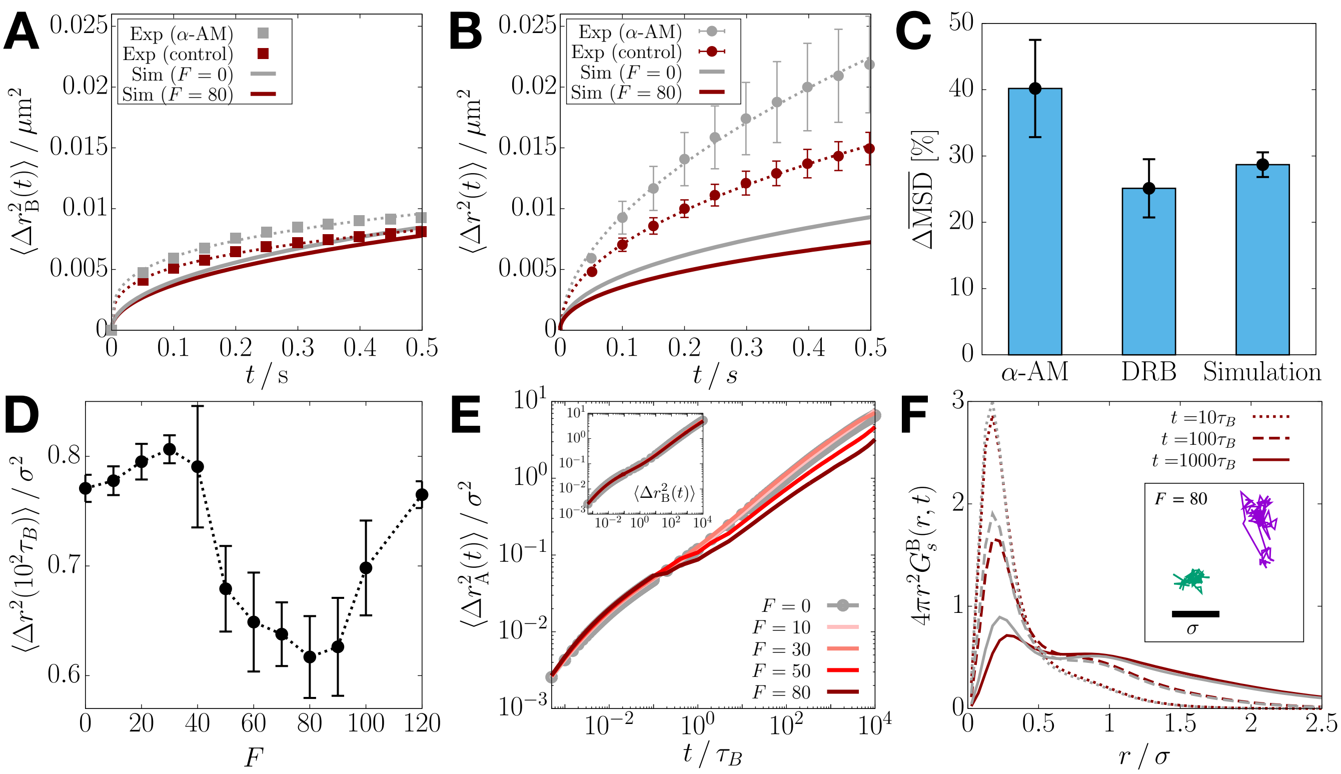


**Fig. S1.** Effect of active forces on the B-type loci dynamics is negligible. (A) Plot of $\langle \Delta r_{\mathrm{B}}^2(t) \rangle$ after treatment with $\alpha$-AM (S4) (squares), which we mimic by turning off the active force (solid lines). Dotted lines are the curves obtained by fitting the experimental data for the control (dark-red) and the inhibited (gray) (B) Same as panel A, except for the plots correspond to overall MSD for all the loci. (C) Bar graphs comparing the increase in the overall MSD (as shown in panel B) relative to the control case when transcription is inhibited using $\alpha$-AM and DRB. Simulations mimic these cases using $F = 0$ and $F = 80$. (D) Overall MSD at $t = 100\tau_B$ as a function of $F$. The dotted lines are a guide to the eye. (E) Log-log plot of the MSDs for the A-type loci with different $F$. The inset shows the same plot for the B-type loci. (F) van Hove functions for the B loci with $F = 0$ (gray) and $F = 80$ (dark-gray) at $t = 10\tau_B$ (dotted line), $100\tau_B$ (dashed line), and $1000\tau_B$ (solid line). The inset shows the 2-D projection of the trajectories of an active (green) and an inactive (purple) loci for $10^4\tau_B$ at $F = 80$.

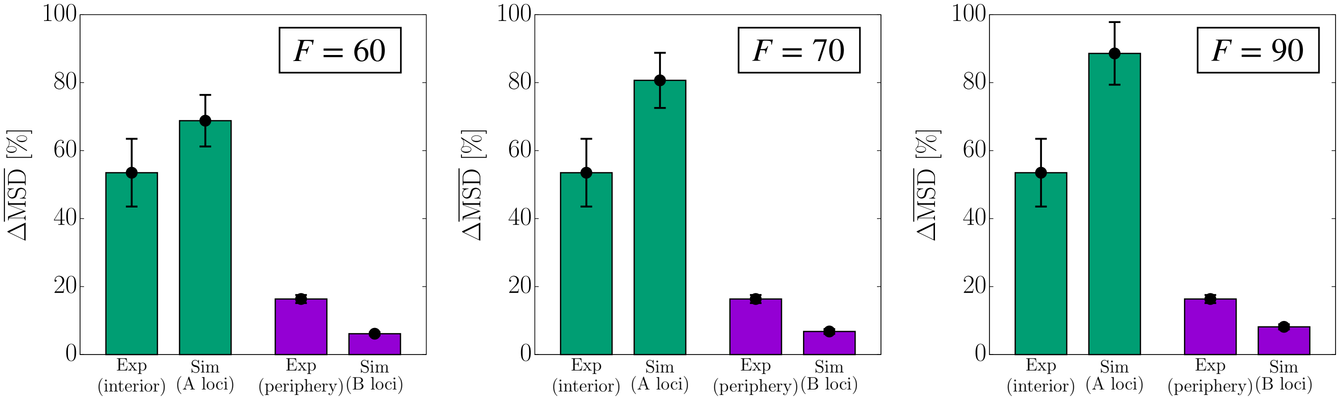


**Fig. S2.** Simulations in the intermediate range of $F$, $60 \leq F \leq 90$, give results that are in a good agreement with experiments. Bar graphs comparing $\Delta\overline{\mathrm{MSD}}$ for the A and B loci separately [Eqs. (S1)-(S2)] between the experiment and simulation results with $F = 60$ (left), 70 (center), and 90 (right). The results for $F = 80$ are given in the main text (Fig. 2B). Taken together, these results show that over the range, $60 \leq F \leq 90$, the agreement with experiments is reasonable, which further reinforces the robustness of the proposed mobility inhibition mechanism.

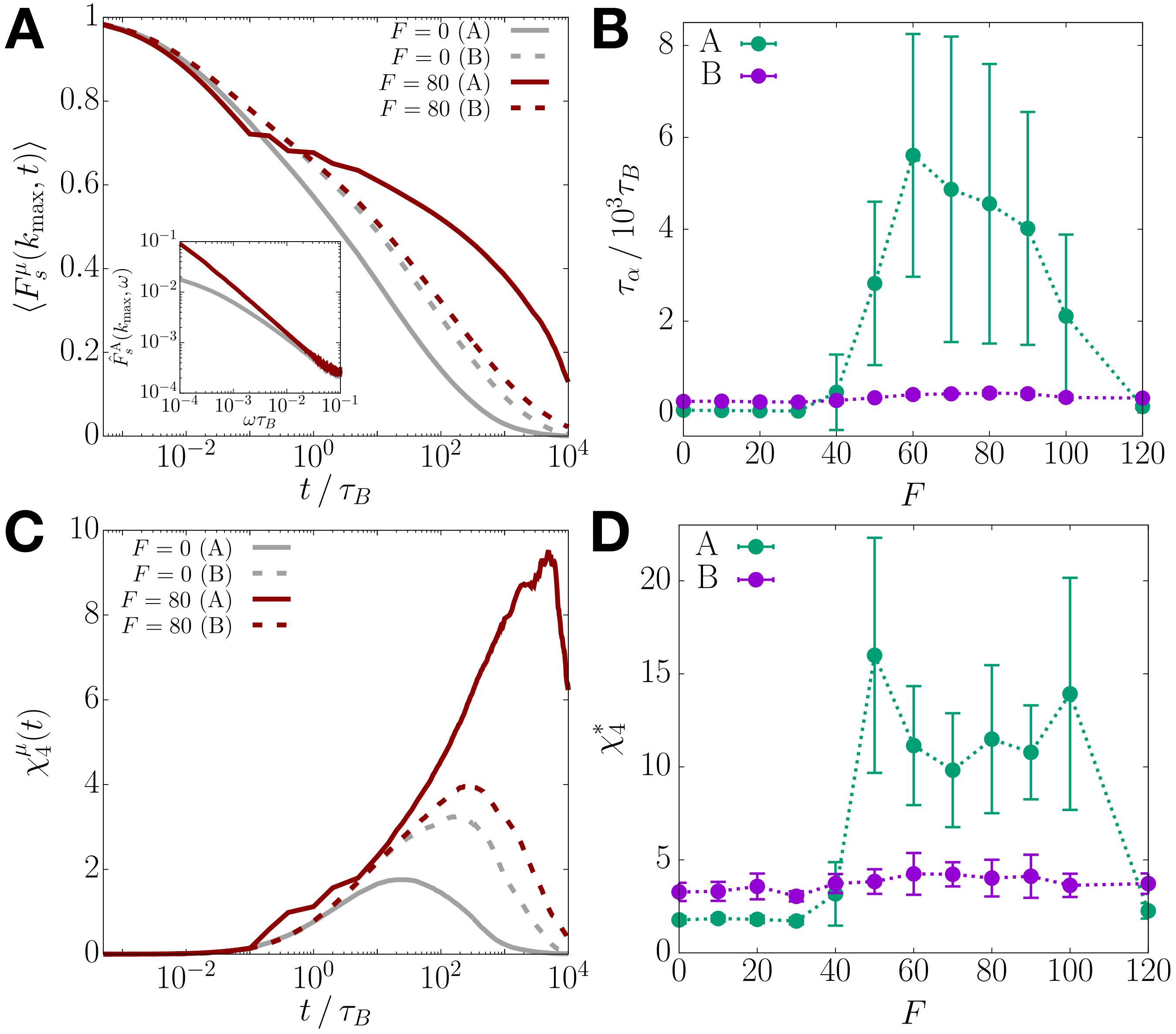


**Fig. S3.** Slow dynamics and heterogeneous mobilities are associated with the euchromatin loci. (A) Comparison between $\langle F_s^{\rm A}(k_{\max}, t)\rangle$ and $\langle F_s^{\rm B}(k_{\max}, t)\rangle$ (Eq. S3), shown by solid and dashed lines, respectively, for $F = 0$ and $F = 80$. The inset shows the log-scale plot of $\hat{F}_s^{\rm A}(k_{\max}, \omega)$, the frequency-domain Fourier transform of $\langle F_s^{\rm A}(k_{\max}, t)\rangle$. (B) The relaxation time of A and B loci for different $F$ values. (C) Comparison between $\chi_4^{\rm A}(t)$ and $\chi_4^{\rm B}(t)$ (Eq. S4), shown by solid and dashed lines, respectively, for $F = 0$ and $F = 80$. (D) The maximum value in $\chi_4^{\rm A}(t)$ and $\chi_4^{\rm B}(t)$ for different $F$. The dotted lines in panels B and D are a guide to the eye.

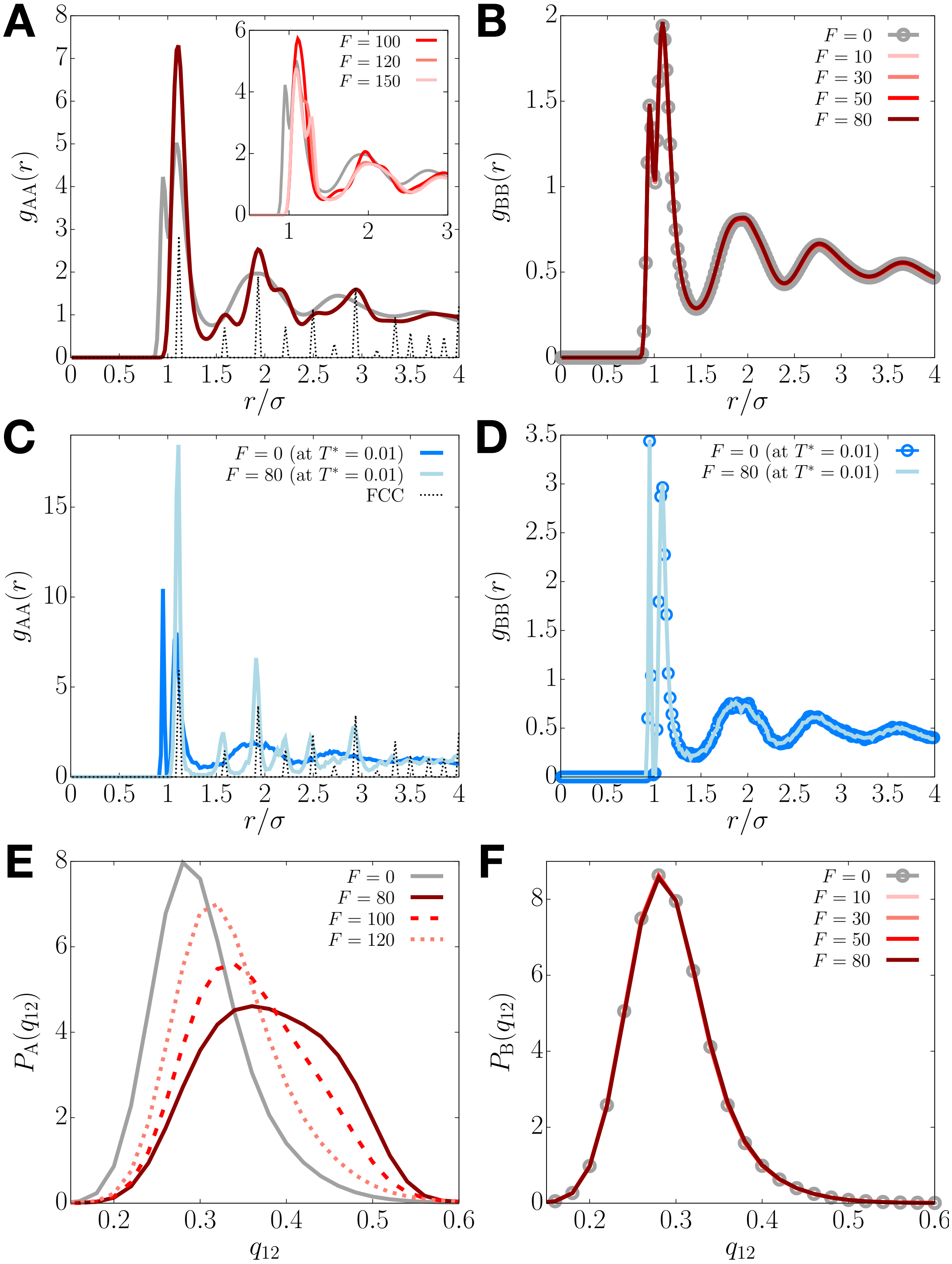


**Fig. S4.** $F$-dependent structural properties for A and B loci. (A) Radial distribution function for A-A loci pairs, $g_{AA}(r)$, at $F = 0$ (gray), $F = 80$ (maroon), and $F > 80$ (inset). $g(r)$ for a FCC crystal is shown with the dotted line for comparison (scaled arbitrarily). The inset shows that the peaks associated with the FCC phase disappear when $F$ exceeds 100. The peak at $r \lesssim \sigma$ for $F = 0$ corresponds to the bonded pairs ($i^{\text{th}}$ and $(i+1)^{\text{th}}$ loci), whose location shifts to the right as $F$ increases. (B) Plots of $g_{BB}(r)$ for different $F$. $g_{BB}(r)$ is independent of $F$ and shows the behavior that is reminiscent of a dense fluid at all $F$ values, including $F = 0$. (C) Inherent structure $g_{AA}(r)$ for $F = 0$ (blue) and $F = 80$ (light blue). (D) Same as panel C except the results for the B-type loci. (E) Distributions of the BOO parameter for the A-type loci [Eqs. (12)-(14)] for $F \geq 80$ and $F = 0$. (F) Same as panel E, except the results are for the B-type loci. Like $g_{BB}(r)$, $P_B(q_{12})$ is independent of $F$.

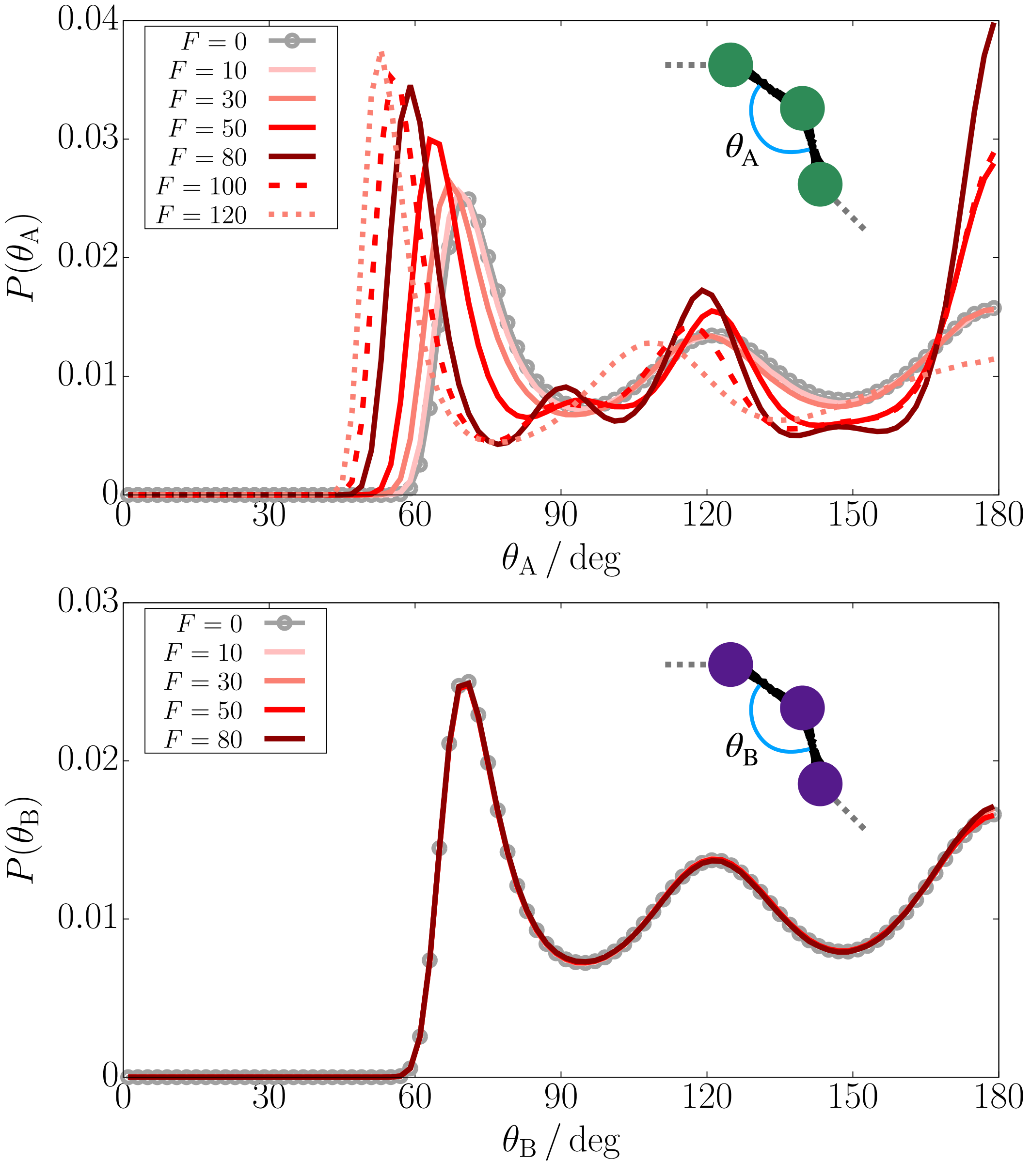


**Fig. S5.** Bending angle distributions for three consecutive A (top) and B (bottom) loci (Eq. S5) at different $F$.

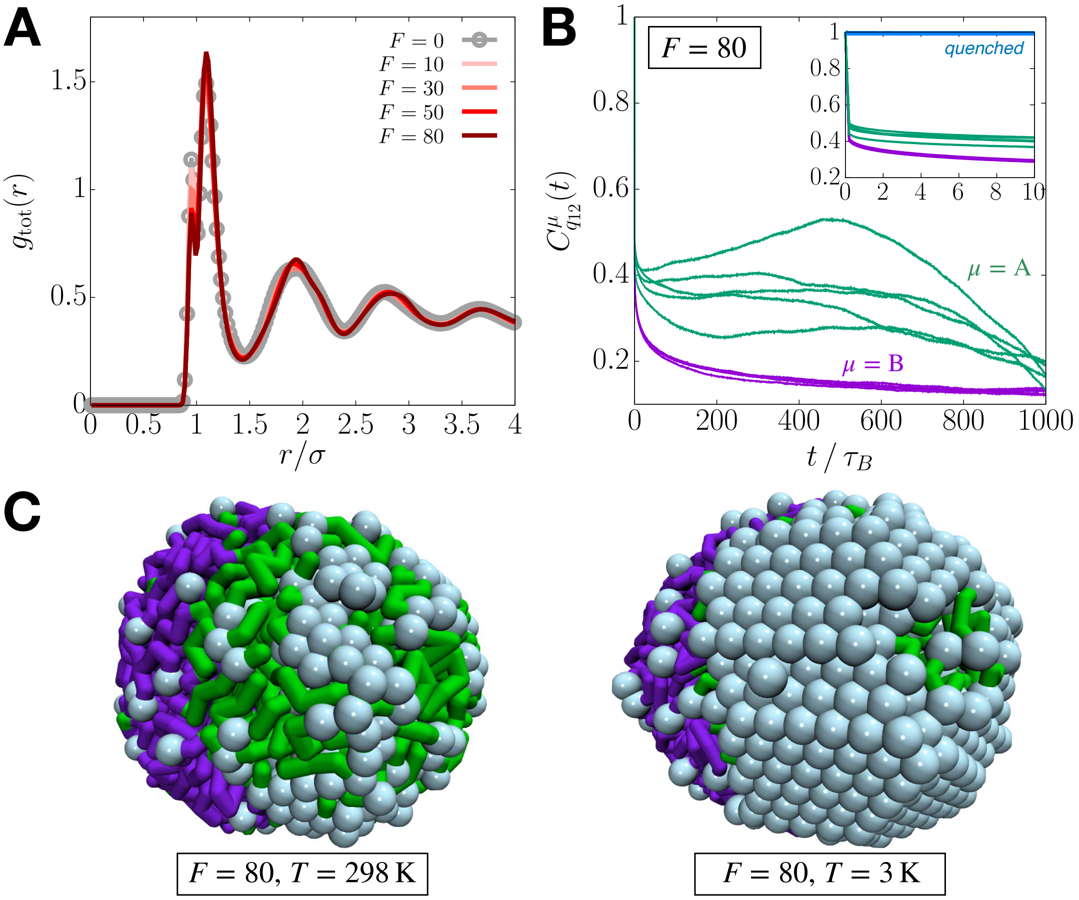


**Fig. S6.** Activity-induced order in the A loci exists only transiently. (A) Radial distribution function for all the locus pairs at different $F$. (B) Plots of the time correlation functions, $C^{\mathrm{A}}_{q_{12}}(t)$ and $C^{\mathrm{B}}_{q_{12}}(t)$ (Eq. S6), for five independent trajectories at $F = 80$. The inset shows the short-time changes in the correlation functions, where $C^{\mathrm{A}}_{q_{12}}(t)$ for the quenched polymer is shown in blue. (C) Simulation snapshots for the active copolymer with $F = 80$ at room temperature (left) and the one quenched to low temperature (right). Green and purple colors represent the A and B loci, respectively. The light blue spheres indicate the loci with $q_{12}(i) > 0.45$. The movies for these simulation trajectories are given in Movies S1 (left) and S2 (right).

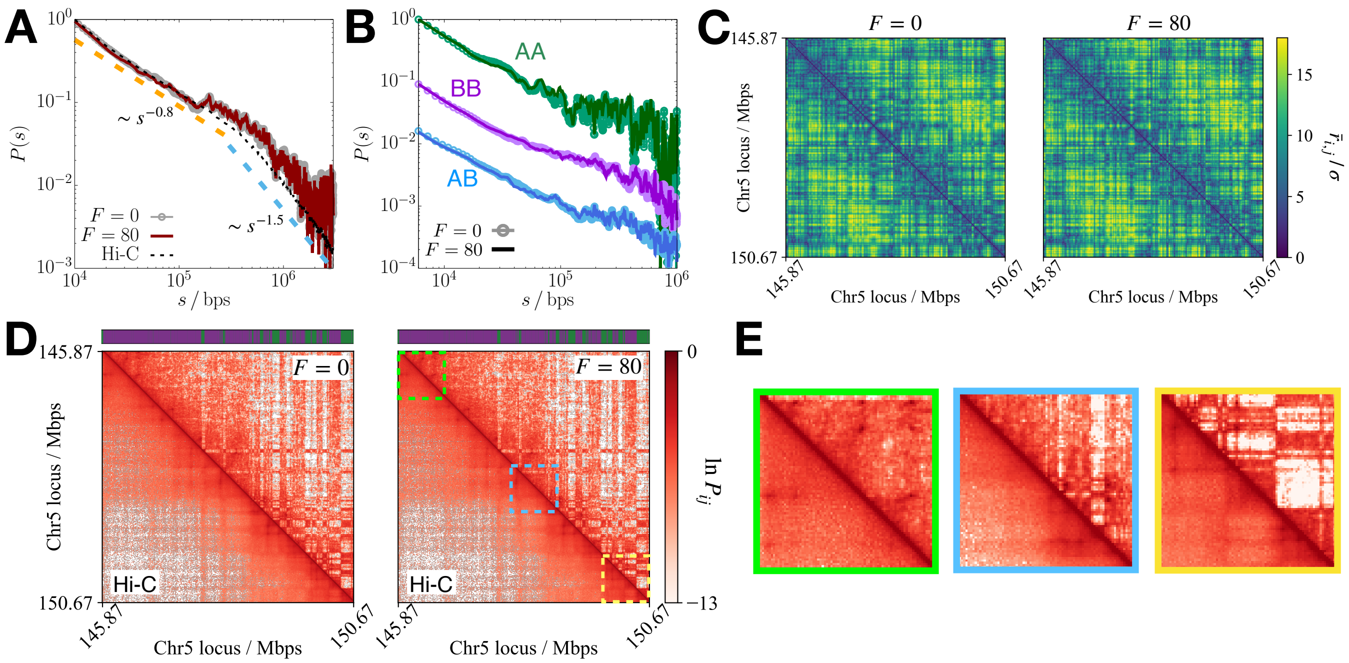


**Fig. S7.** Solid-like ordering does not alter the chromatin conformation. (A) Plot of $P(s)$ (Eq. S7) at $F = 0$ (gray) and $F = 80$ (dark red). For comparison, $P(s)$, calculated from the Hi-C data (S12), is shown in black dashed line. Two distinct power-law decays are indicated by the orange and blue dashed lines. Each $P(s)$ curve is normalized such that it decays from unity. (B) Plots of $P_{\mu\gamma}(s)$ (Eq. S8) at $F = 0$ (circles) and $F = 80$ (solid line) for A-A (green), B-B (purple), and A-B (blue) pairs. $P_{\text{AA}}(s)$ is normalized such that it decays from unity, whereas $P_{\text{BB}}(s)$ and $P_{\text{AB}}(s)$ are shifted from $P_{\text{AA}}(s)$ in the negative y-direction for a better visualization. (C) Heat map for the mean distance matrices obtained from the single trajectories at $F = 0$ (left) and $F = 80$ (right), which start from an identical configuration. (D) Comparison between the simulated contact maps (upper triangle) and the Hi-C contact map (lower triangle), where the A/B (green/purple) profile in the model is shown on the top of each map. The predicted contact maps at $F = 0$ (left) and $F = 80$ (right) are essentially the same. (E) Close-up views of three 0.8-Mb regions highlighted by the green, blue, and yellow dashed squares in the righthand contact map in panel D. The green and blue regions show good agreement between the simulated and Hi-C contact maps, whereas there are some quantitative deviations in the yellow region. Such deviations could arise from the simplicity of the model as well as the finite-size effects, which are in the yellow region that includes a terminus of the polymer with the alternation of locus type. Note that the green region at the other end, which consists of predominantly B-type locus, shows excellent agreement. The blue region in the middle of the polymer also shows good agreement. Further adjustment of the interaction parameters ($\epsilon_{\text{AA}}$, $\epsilon_{\text{BB}}$, $\epsilon_{\text{AB}}$) would yield a better agreement, which we did not carry out here.

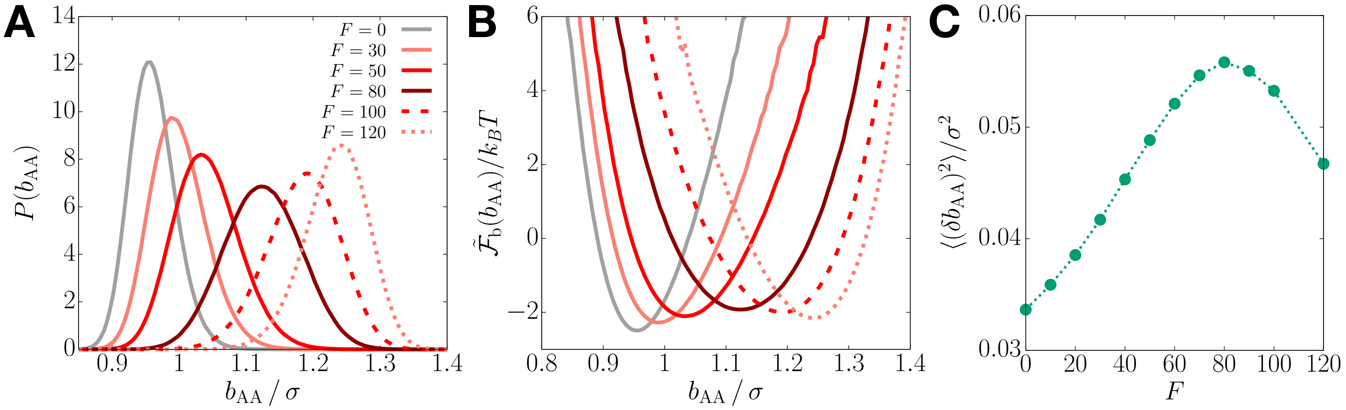


**Fig. S8.** Explanation of the ordering mechanism using activity-induced bond length extension. (A) Probability distribution of the A-A bond distance for different $F$ values. (B) Plots of the effective bond free energy defined by $\tilde{\mathcal{F}}_{\mathrm{b}}(b_{\mathrm{AA}}) = -k_BT \ln P(b_{\mathrm{AA}})$. (C) Bond length fluctuations as a function of $F$. The dotted lines are a guide to the eye.

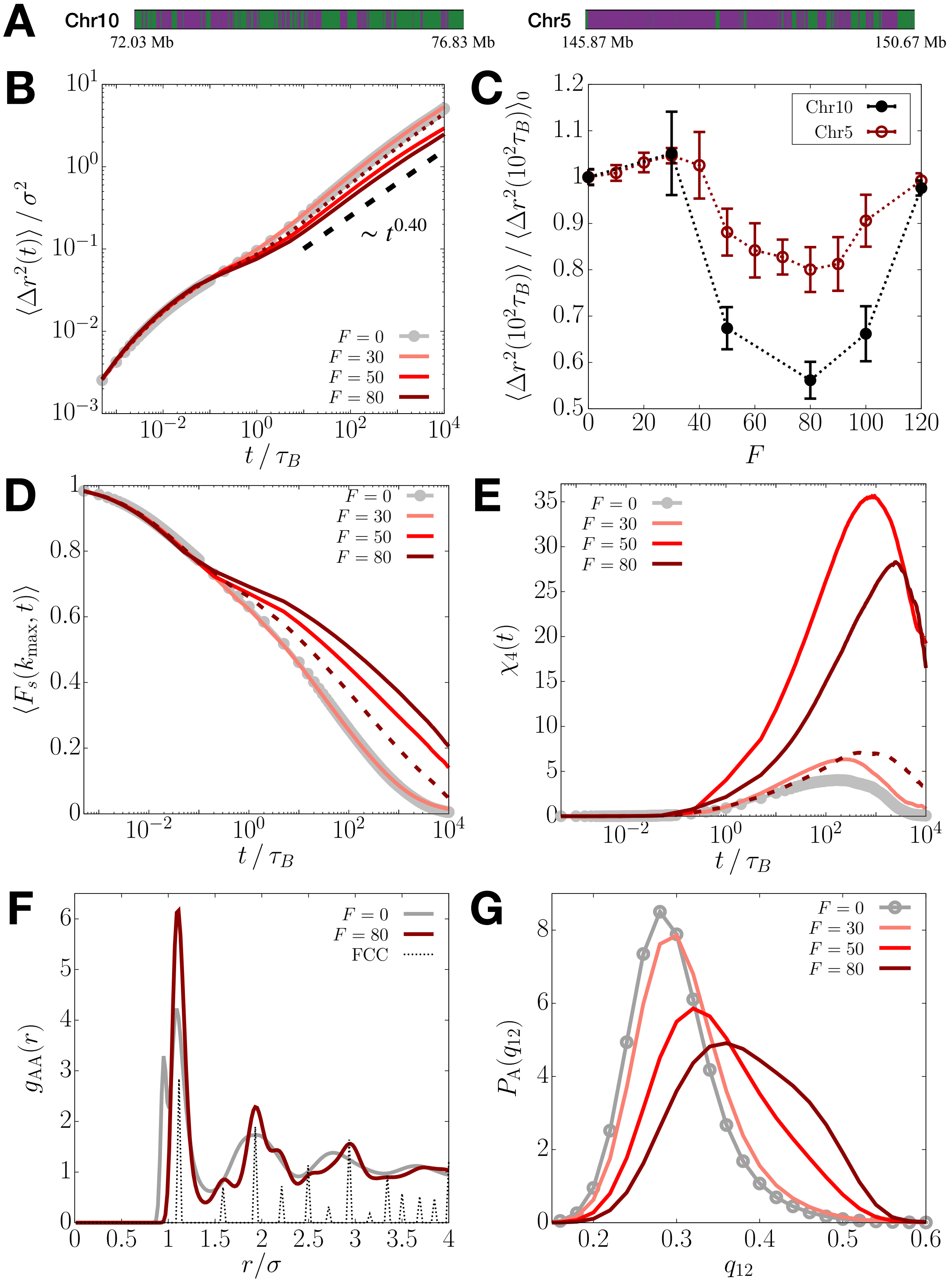


**Fig. S9.** Dynamical properties for a 4.8-Mbp segment of Chromosome 10 (Chr10). (A) Comparison between the A/B (green/purple) profiles for the Chr10 (left; $N_A/N = 0.58$) and Chr5 (right; $N_A/N = 0.25$) segments. (B) Log-log plots of MSDs for different values of the activity, $F$. For comparison, the MSD curve for Chr5 with $F = 80$ is shown by the dotted line. (C) Ratio of the MSD at $t = 100\tau_B$ for a given activity level $F$ to the passive case, where the data for Chr10 (black) and Chr5 (dark-red) are compared. The dotted lines are a guide to the eye. (D and E) Plots of $\langle F_s(k_{\max}, t) \rangle$ and $\chi_4(t)$ for different $F$ values. The data for Chr5 with $F = 80$ are shown by the dashed lines. (F) $g_{AA}(r)$ at $F = 0$ (gray) and $F = 80$ (maroon). $g(r)$ for a FCC crystal is shown with the dotted line for comparison. (G) Plots of $P_A(q_{12})$ at different $F$.

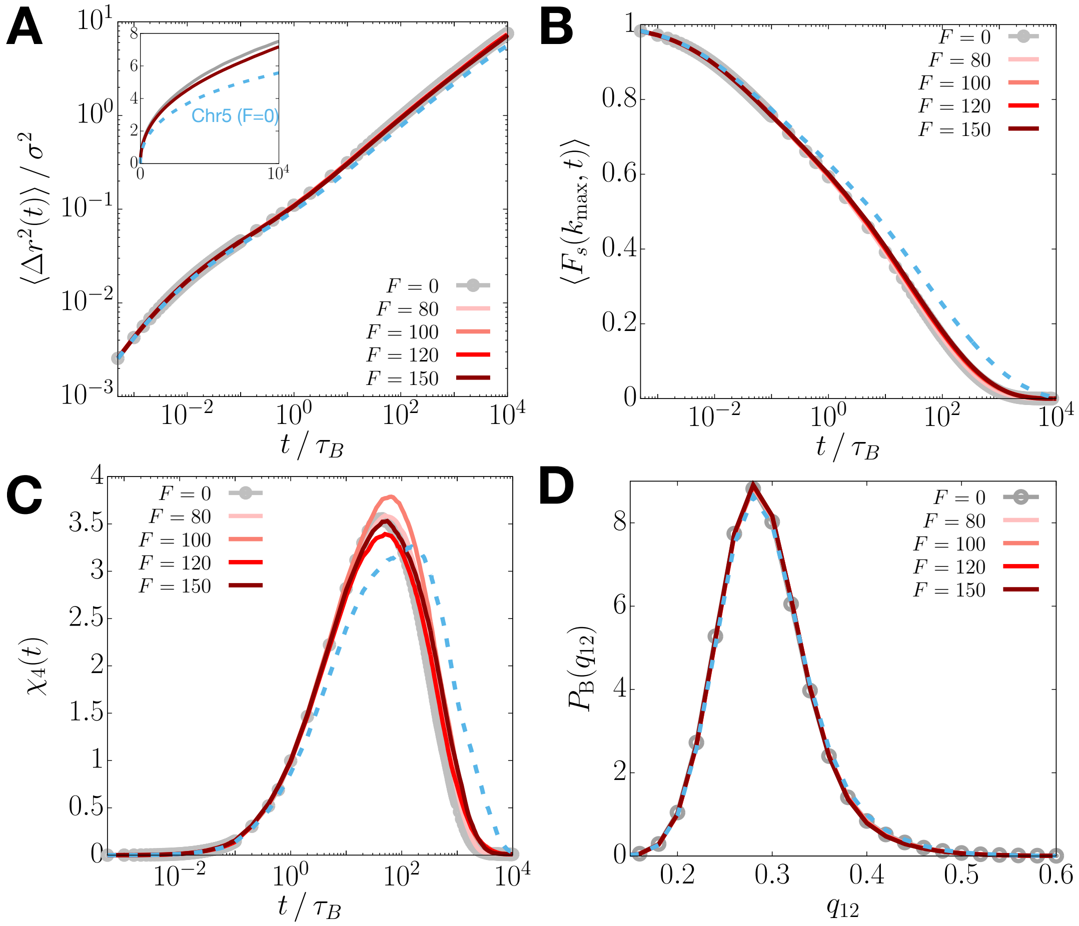


**Fig. S10.** A copolymer chain with the same length and the fraction of active loci as Chr5 but randomly shuffled epigenetic sequence does not show dynamical changes and structural ordering upon increasing $F$. (A) Log-log plots of the MSDs for different values of activity, $F$. For comparison, the MSD curve for the passive Chr 5 is shown by the dashed line. The inset shows the plots in regular scale. (B and C) Plots of $\langle F_s(k_{\text{max}}, t) \rangle$ and $\chi_4(t)$ for different $F$ values. The data for the passive Chr5 are shown by the dashed lines. (D) Probability distributions of $q_{12}$ for B loci at different activity levels, where the dashed line is the distribution for Chr5 with $F = 0$, respectively.

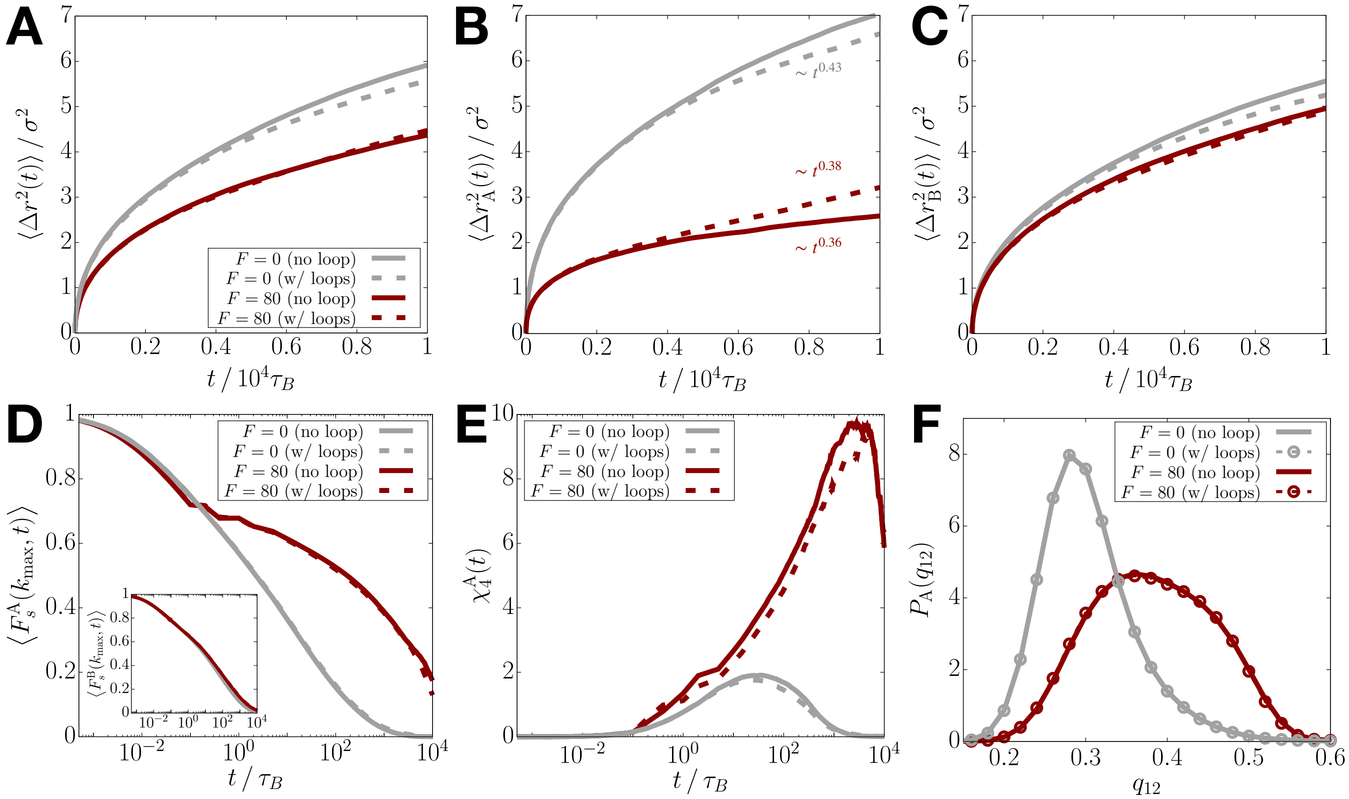


**Fig. S11.** Removing the CTCF-mediated loops does not alter the activity-induced suppressed mobilities. (A) MSD as a function of time for the Chr5-CCM at $F = 0$ (gray) and $F = 80$ (dark-red), with (dashed) or without (solid) the CTCF-mediated loops. (B and C) Same as panel A, except the plots show $\langle \Delta r_{\mathrm{A}}^2(t) \rangle$ and $\langle \Delta r_{\mathrm{B}}^2(t) \rangle$. (D) Plots of $\langle F_s^{\mathrm{A}}(k_{\mathrm{max}}, t) \rangle$ and $\langle F_s^{\mathrm{B}}(k_{\mathrm{max}}, t) \rangle$ (inset) (Eq. 3) at $F = 0$ (gray) and $F = 80$ (dark-red), with (dashed) or without (solid) the loops. (E) Same as panel D, except showing the plots of $\chi_4^{\mathrm{A}}(t)$ (Eq. 4). (F) Probability distribution of the BOO parameter for the A loci at $F = 0$ (gray) and $F = 80$ (dark-red), with (dashed) or without (solid) the loops.

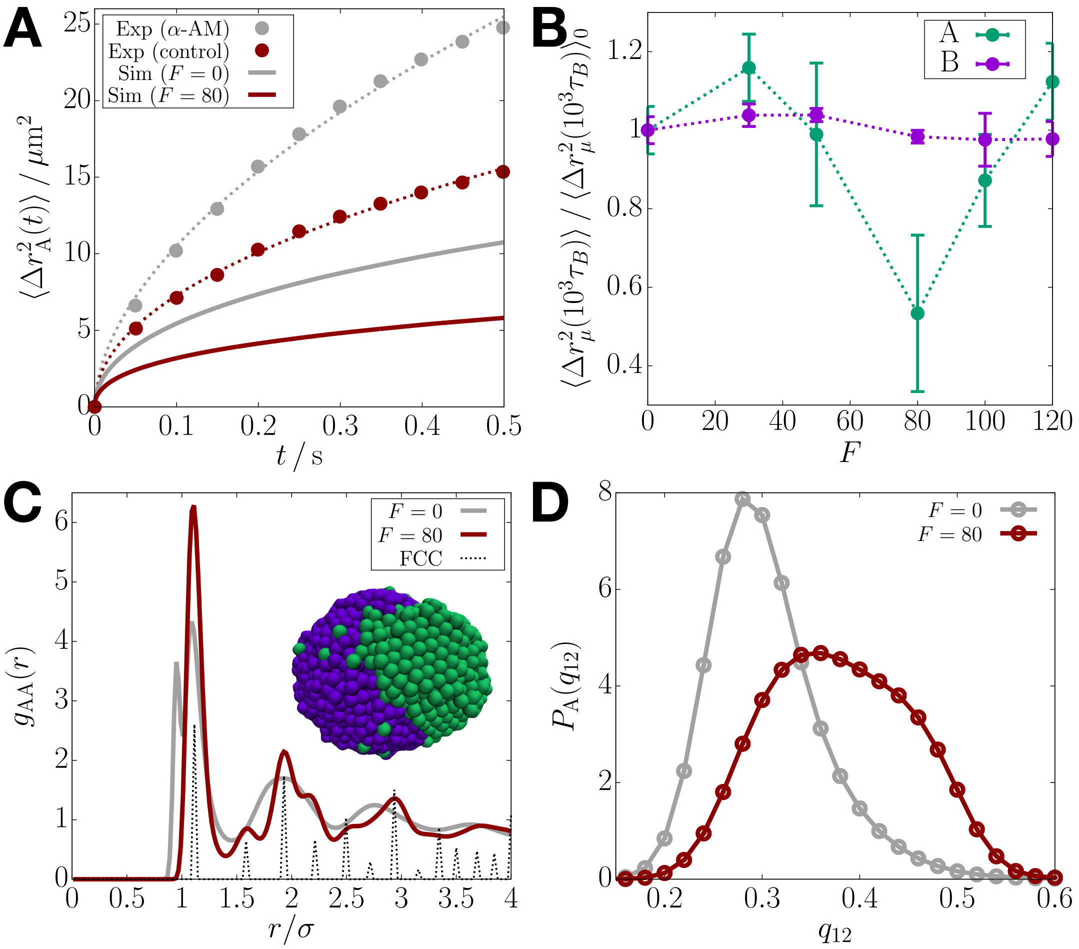


**Fig. S12.** Choice of the non-bonding interaction parameters ($\epsilon_{AA} < \epsilon_{BB}$) does not alter the results of the activity-induced suppressed mobilities and transient ordering. (A) Plots of $\langle \Delta r_A^2(t) \rangle$ from the CCM simulations for Chr5 with $F = 0$ and $F = 80$ (solid lines), compared with the euchromatin MSD from the experiment that inhibits transcription using $\alpha$-AM (S4) (circles). (B) Ratio of the MSD at $t = 10^3 \tau_B$ for a given activity level $F$ to the passive case, where the data for A (green) and B (purple) loci are compared. The dotted lines are a guide to the eye. (C) $g_{AA}(r)$ at $F = 0$ (gray) and $F = 80$ (maroon). $g(r)$ for a FCC crystal is shown with the dotted line for comparison. A simulation snapshot for $F = 80$ is shown. (D) Plots of $P_A(q_{12})$ at $F = 0$ (gray) and $F = 80$ (maroon).

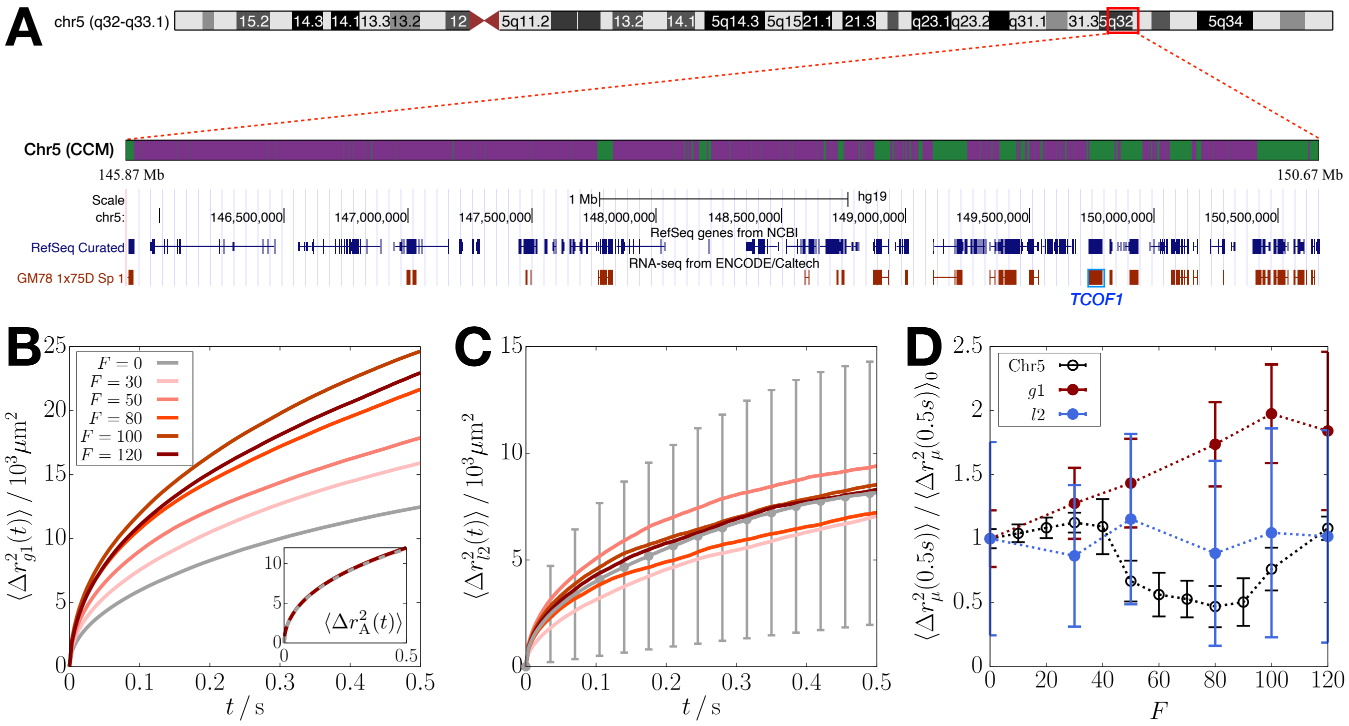

**Fig. S13.** Application of the active forces to only one gene region or a few loci enhances mobility. (A) Screenshot from the UCSC Genome Browser (http://genome.ucsc.edu) (S13) for the region of our interest. The top panel shows the ideogram of Chr5. The region modeled using the CCM (Chr5: 145.87–150.67 Mb) is highlighted in the red box. In the bottom panel, the first track shows the A/B (green/purple) profile in the CCM. The second track (navy color; right below the scale bar) shows the locations of all the genes in the given region, and the third one (maroon color) indicates the active genes in the GM12878 cell line. The *TCOF1* gene is marked in blue. (B) Time-dependent MSD for the loci corresponding to the *TCOF1* gene body (labeled "*g1*") as a function of $F$. The inset shows the MSD for all the A loci at different values of $F$. (C) $F$-dependence of the MSD for the first two consecutive A-A bonds in the *TCOF1* gene region (labeled "*l2*"). The color scheme is the same as that for panel B. (D) Ratio of the MSD at $t = 0.5\,\mathrm{s}$ at $F$ to the passive case ($F = 0$). The data for the single active gene (dark-red) and the two A-A bonds (blue) are compared with the results for the Chr5 region in panel A with all the genes activated (black; *cf.* Fig. S1D). The dotted lines are a guide to the eye.

**Movie S1. A simulation trajectory for Chr5 at $F = 80$ and $T = 298\,\mathrm{K}$, corresponding to $2500\tau_{\mathbf{B}} \approx 1.75\,\mathrm{s}$. Green and purple colors represent the A and B loci, respectively. The light blue spheres indicate the loci with $q_{12}(i) > 0.45$.**

**Movie S2. A simulation trajectory for Chr5 at $F = 80$ and $T = 3\,\mathrm{K}$ (quenched). The color scheme is the same as in Movie S1.**
